# Good and bad lettuce leaf microbes? Unravelling the genetic architecture of the microbiome to inform plant breeding for enhanced food safety and reduced food waste

**DOI:** 10.1101/2021.08.06.455490

**Authors:** A Damerum, EC Arnold, V Bernard, HK Smith, G Taylor

## Abstract

Lettuce is a high value food crop, consumed raw around the world. Engineering of the leaf microbiome could provide significant benefits for enhanced crop yield and stress resistance and help to reduce food waste caused by microbial spoilage. Lettuce leaves also act as a vector for human pathogens, implicated in several high-profile food-borne disease outbreaks. Since host genotype helps determine microbiome composition, we hypothesize that leaf surface traits can be defined that associate with ‘good’ bacterial microbiomes providing benefits to the crop and that ‘bad’ microbiomes, where spoilage organisms and human pathogens are abundant, can also be associated to underlying leaf genetics, providing key targets for future crop breeding. Using a Recombinant Inbred Line (RIL) population, we show that cultivated and wild parental genotypes differ with reduced bacterial diversity, larger leaves and fewer, larger stomata, smaller epidermal cells and more hydrophilic leaf surfaces found in the cultivated compared to wild lettuce. Functional analysis of the associated microbiome revealed increased abundance of genes associated with disease virulence for the cultivated lettuce genotype, suggesting domestication has had broad impacts on leaf and associated bacterial microbiome traits. We defined the core lettuce bacterial microbiome from 171 RILs, comprised of 45 taxa in the phyla Proteobacteria, Actinobacteria, Firmicutes, Chloroflexi and Deinococcus-Thermus. Leaf surface characteristics important in influencing bacterial diversity and abundance were identified as stomatal size (length and width), epidermal cell area and number and leaf surface hydrophobicity of the abaxial leaf surface. Quantitative trait loci (QTL) for leaf surface traits, frequently mapped alongside those for the extended phenotype of bacterial taxa abundance, including for human pathogens *Campylobacter* spp., *Escherichia-Shigella* spp., *Clostridium* spp. (LG 4, 5 and 6) and spoilage bacteria, *Pseudomonas* spp. (LG 1, 3, 4, 6 and 9). Candidate genes underlying these QTL were enriched in GO terms for cell wall assembly and modification, defence response, hormone-mediated signalling and biosynthesis and anatomical structure development. This work provides the first insight into the genetic architecture of host surface traits in a leafy crop alongside the mapped genetic architecture of bacterial communities and has identified areas of the lettuce genome as important targets for future microbiome engineering.

## INTRODUCTION

The phyllosphere microbiome, including all microorganisms associated with the aerial portions of plants, is a dynamic ecosystem, comprised primarily of bacteria and fungi and governed by host genetics, environmental conditions and microbe-microbe interactions. The coevolution of land plants and microbes over ~400 million years has forged complex relationships, which can be mutualistically beneficial, commensal or pathogenic and have important implications for plant health and productivity (Laforest-Lapointe *et al.*, 2017). Imbalances in the phyllosphere microbiome composition, termed phyllosphere dysbiosis (Liu *et al.*, 2020), can impact crop health pre-harvest, translating into yield and crop losses, can increase rates of spoilage, reducing shelf life post-harvest, leading to food waste and can have serious implications for food safety.

Lettuce (*Lactuca sativa* L) is the most widely consumed leafy vegetable, with the US market valued at $3.5 billion in 2019 (USDA, 2020). The lettuce phyllosphere microbiome is dominated by bacteria, with 10^5^ to 10^7^ bacterial cells per g of leaf tissue, the majority of which are commensal (Williams *et al.*, 2013), with a small number of representative phyla including Proteobacteria, Firmicutes, Actinobacteria and Bacteroidetes accounting for the majority of microbiota present, although overall proportions can vary seasonally, temporally (even daily) and with differing irrigation regimes (Williams *et al.*, 2013). Post-harvest, cold storage conditions can accelerate the growth of psychrotrophic bacteria, such as *Pseudomonas* and *Erwinia*, which produce pectinolytic enzymes resulting in soft rot, deteriorating both organoleptic and textural properties (Ragaert *et al.*, 2007), shortening shelf-life and leading to spoilage.

The lettuce phyllosphere is important for human health, as a vegetable which is largely consumed without cooking. Leafy vegetables are now known to be the biggest cause of foodborne illness in the US, resulting in 47 deaths from 1998-2013 (Herman *et al.*, 2015; Bennett *et al.*, 2018). Human pathogenic bacteria including *Salmonella* spp., *Escherichia coli* O157:H7, *Shigella* spp., *Clostridium* spp., *Campylobacter* spp. and *Listeria monocytogenes* are the leading causes of foodborne illness in raw produce (Bennett *et al.*, 2018) and are difficult to detect, given that they have no observable impact on crop health. Sources of contamination include pre-and post-harvest include soil, irrigation and groundwater, manure, animal faeces (predominantly livestock) and factory worker handling (Alegbeleye *et al.*, 2018). Currently, washing leaves with chemical disinfectants in an attempt to achieve surface sterilisation is used for decontamination, but this has limited efficacy. Following chlorine treatment, food-borne pathogens *L. monocytogenes* and *Salmonella enterica* have been shown to enter a viable but non-culturable state, enabling them to survive the stress and not be detected by culture-based methods, but remain virulent (Highmore *et al.*, 2018). Washing is also ineffective at irradicating internalised pathogens, or those which contaminate leaves post-washing.

The leaf epidermis provides the first physical barrier to microbe colonisation and is the site of initial pathogen recognition and subsequent defence responses. Leaf surface characteristics such as vein and stomatal density, hydrophobicity and soluble protein concentrations are known to influence attachment of *Salmonella enterica* on lettuce leaves (Hunter *et al.*, 2015). Bacterial cells can also aggregate in epidermal cell junctions and stomatal pores provide entry routes into internal leaf spaces (López-Gálvez *et al.*, 2010). Cells of *E. coli* preferentially reside within stomatal pores, which have provided shelter, even after washing (López-Gálvez *et al.*, 2010). Whilst plant detection of *E. coli* O157:H7 can trigger stomatal closure to prevent internalisation, the plant pathogen *Pseudomonas syringae* has been shown to override this immune response, reversing stomatal closure (Zeng *et al.*, 2010). Minimal processing of lettuce including washing and drying, which is common practice in lettuce, can result in bruising of leaf surfaces, increasing *E. coli* O157:H7 population sizes (Brandl, 2008). Interestingly, leaves already infected by soft rot or displaying abiotic stress-induced tipburn show increased colonisation of *E. coli* O157:H7 compared to healthy leaves (Brandl, 2008), highlighting the complex interplay of not only host-microbe, but microbe-microbe interactions in influencing phyllosphere microbiome dynamics.

Plant-associated microbiomes are known to confer fitness advantages to the plant host (Trivedi *et al.*, 2020) but despite the benefits of a functioning ‘holobiont’, where evolutionary adaptations of host and microbes act together to determine stability, few studies have identified the genetic basis of plant traits that could be influencing the composition of the associated microbiome and the genetic architecture of the microbiome. Genetic mapping of abundance, diversity and composition has rarely been reported, despite the power and impact that such data could have on future plant breeding. For lettuce, there is some indication that plant genotype can influence the attachment and proliferation of human pathogenic bacteria (Hunter *et al.*, 2015; Jacob & Melotto, 2020), with differences in the sequenced meta-barcoded microbiome also apparent across contrasting lettuce genotypes (Hunter *et al*., 2010). Whilst investigations have generally focused on the interaction of one or few strains with the host, unravelling complex host-microbe interactions requires investigation of community assemblage as a whole. Genome-wide association studies are beginning to address this gap, but remain largely focussed on limited microbial taxa (Beilsmith *et al.*, 2019). Identifying host genes which influence both bacterial community structure and specific taxa could enable suitable ‘good’ microbiome selection through plant breeding and indirect microbiome engineering. Additionally, elucidating microbe-microbe interactions may enable the development of probiotics, applied in agricultural systems to outcompete ‘bad’ pathogenic bacteria and improve plant health (Koskella, 2020).

The aim of this study was to address this important knowledge gap on the genetic architecture of the extended phenotype between host and microbiome. Using 16S rRNA metabarcoding of the bacterial phyllosphere communities of a lettuce Recombinant Inbred Line (RIL) mapping population, we mapped QTL for leaf surface characteristics, including stomata and epidermal cell patterning and cuticle hydrophobicity, alongside QTL for bacterial species richness and abundance, providing new insight into the genetic basis of this extended phenotype of high relevance to food spoilage and safety. We identified host traits and genomic regions governing bacterial species richness and abundance, along with underlying candidate genes and discuss how future lettuce breeding can be deployed to positively manipulate the phyllosphere microbiome.

## METHODS

### Plant cultivation, harvesting and phenotyping

A total of 205 recombinant inbred lines (RILs) generated from a cross between *Lactuca sativa* cv. Salinas × *Lactuca serriola* accession US96UC23 (Truco *et al.*, 2007, 2013) were grown on a commercial farm near Azenha do Mar, Portugal (37°28′25.2“N; 8°47′42.2“W) in Spring 2016. Seedlings were sown 10 cm apart in three 1.2 × 35 m beds, each representing a separate block, in a randomised block design with three replicates of each RIL and a row of border plants surrounding each bed, as described previously (Zhang *et al.*, 2007). Plants were watered by sprinkler irrigation according to commercial practice.

For leaf surface morphology traits, the fourth true leaf was harvested from each plant 39 d after sewing, refrigerated and processed within 24 h of harvest. The leaf was imaged using a EOS 300D camera (Cannon, UK) to estimate leaf area (LA; mm^2^) and abaxial and adaxial epidermal imprints were collected as described by Zhang *et al.* (2007), which were later imaged using an Axioplan2 light microscope (Zeiss, UK), to obtain measurements of epidermal cell area (ECA; μm^2^), cell number (ECN; count per mm^2^), stomatal density (SD; per mm^2^), stomata width (SW; from outer guard cell pair, μm) and length (SL; μm). Stomatal index (SI; %), a measure of the proportion of epidermal cells formed as stomatal cells was calculated a described previously (Zhang *et al.*, 2007). Leaf (LC) and epidermal cell (ECC) circularity index were calculated using the following formula:

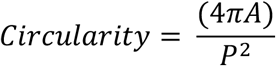

Where *A* is leaf or cell area and *P* is leaf or cell perimeter, with an index of 1 for a perfect circle. Leaf contact angle (CA, θ) was measured by dropping 10 μL of distilled water onto the centre of a 1 cm^2^ leaf strip, imaging the droplet with a EOS 500D camera fitted with a Cannon EF 50mm f/2.5 compact macro lens (Cannon, UK) and measuring the angle tangent to the water droplet (Lamour *et al.*, 2010). All images were analysed using ImageJ version 1.50e (Schneider *et al.*, 2012), using the ‘multi-point’, ‘straight-line’, ‘area’ and ‘angle’ tools.

For 16S rRNA sequencing, five leaves were aseptically harvested from 171 of the RILs in triplicate, flash-frozen in liquid nitrogen and stored immediately on dry ice before longer-term storage at −80 °C, until nucleic acid extraction.

### Bacterial community profiling and analyses

In order to isolate external and internalised bacteria, whole leaves were aseptically ground in liquid N_2_ in a pestle and mortar. DNA was extracted from ~150 mg of leaf tissue using a CTAB protocol (Doyle & Doyle, 1987) followed by ethanol precipitation. Three replicates of 171 RILs (one per experimental block) and six replicates of each RIL population parent (two per experimental block) totalling 525 samples were analysed by 16S rRNA profiling using the chloroplast excluding 799F (5’-AACMGGATTAGATACCCKG-3’) and 1115R (5’-AGGGTTGCGCTCGTTRC-3’) primer pair, targeting the V5-V6 region (Kembel *et al.*, 2014). Amplicon libraries were prepared for sequencing using a two-step PCR protocol by QB3 Genomics (UC, Berkeley) and multiplexed libraries were sequenced on three lanes of Illumina MiSeq v3 300 bp paired-end sequencing at the Vincent J. Coates Genomics Sequencing Laboratory (UC, Berkeley, U.S.). Sequences were demultiplexed and converted to FASTQ files using bcl2fastq (v2.19) by the sequencing centre.

A total of 31.7 million reads were obtained from three lanes of MiSeq v3 PE 300 bp sequencing, with a minimum of 22,806 reads and average of 59,701 reads per sample before filtering. 16S rRNA profiling was achieved using the QIIME2 version 2020.2 pipeline and available plugins (Bolyen *et al.*, 2019). Sequences were trimmed, low quality sequences (quality score <30) were eliminated (on average, 82.69% of original reads retained), amplicon sequencing errors were corrected, paired reads were concatenated (74.17% of original reads retained) and chimeric sequences and singletons were removed (71.20% of original reads retained) using DADA2 (Callahan *et al.*, 2016), truncating forward reads to 240 bp and reverse to 220 bp. The resultant amplicon sequence variants (ASVs) were clustered at 99% identity by open-reference clustering using vsearch (Rognes *et al.*, 2016). Taxonomy was assigned by machine-learning-based classification using the q2-feature-classifier classify-sklearn plugin (Bokulich *et al.*, 2018), with the Naïve Bayes classifier trained on full-length 16S rRNA sequences using the SILVA 132 16S rRNA database (Glöckner *et al.*, 2017). The resultant ASV table was filtered to exclude plastid, cyanobacteria and archaea sequences. Approximately 50% of reads were excluded after taxonomy assignment due to rRNA mitochondria contamination, with an average of 22,538 reads per sample remaining. A phylogenetic tree was constructed using an approximately-maximum-likelihood method with FastTree (Price *et al.*, 2010). For the RIL data, the ASV table was filtered to exclude RIL parent samples, the three replicates of each RIL were grouped and the mean-ceiling was calculated before further analyses. ASVs were named as the highest level of classification available, followed by ascending rank position according to total abundance.

### Statistical analyses

To compare significant differences in leaf surface characteristics between parents or extreme RILs (RILs demonstrating extreme high or low trait values for a given trait), either a two-sample t-test or non-parametric Wilcoxon signed-rank test for non-normal data were used, all calculated in R (R Core Team, 2017).

Rarefaction of the two ASV tables comprising the RIL dataset and the RIL parents was performed to the minimum sequence depth, which was 6,845 and 14,176, respectively. All further analyses were conducted on the rarefied dataset unless otherwise stated. Alpha (α; within-sample diversity) and beta (β; between-sample diversity) diversity metrics were calculated using the q2-diversity plugin and the core-metrics-phylogenetic script implemented in QIIME2 (Bolyen *et al.*, 2019) and taxa plots were drawn using the phyloseq package in R (Mcmurdie & Holmes, 2013). For the RIL parents, statistically significant differences in α diversity metrics were calculated using unpaired two-sample t-tests calculated in R (R Core Team, 2017) and differences in multivariate β diversity metrics were calculated using permutational multivariate analysis of variance (PERMANOVA) with 999 permutations in QIIME2. For the RILs, PERMANOVA was used to investigate the partitioning of distance matrices among leaf traits was conducted using the “adonis” function in the vegan package (Oksanen *et al.*, 2020).

Differential abundance analyses of ASVs were performed on non-rarefied data using the phyloseq and DESeq2 packages in R (Love *et al.*, 2014) to compare 1) the wild and cultivated parents and 2) the RILs with extreme high and low trait values (within the top 20 upper and lower tails of trait frequency distribution). In both instances, ASVs tables were first filtered to exclude those present in <5% of the sample set. Functional predictions of ASVs were assigned using PICRUSt2 (Douglas *et al.*, 2020) according to the KEGG Orthology (KO) database, implemented using the QIIME2 plugin, q2-picrust2 (QIIME2 version 2019.10), which were also examined for differential abundance with DESeq2.

Correlation between traits and the top 20 most abundant ASVs was calculated by Spearman’s rank-order correlation coefficient, visualised using the corrplot package in R (Wei & Simko, 2021). Principal component analysis was performed on the ASVs present in >5% RILs and the function *envfit* was used to identify leaf surface morphology traits which had a significant influence, based on a permutation test with 999 iterations using the vegan package in R (Oksanen *et al.*, 2020).

Co-occurrence network analyses were used to investigate interactions between ASVs present in >5% of the RILs. The network was constructed using SPIEC-EASI (Kurtz *et al.*, 2015), which infers an inverse covariance matrix with data transformations developed for compositional data, using the Meinshausen and Buhlmann network method, with 100 repetitions. Network analyses were conducted using igraph in R (Csardi & Nepusz, 2006), including determining the number of adjacent edges (*degree*), average distance between vertices (*mean_distance*), clustering coefficient (*transitivity*), detection of clusters and modularity estimation using the Louvain algorithm (*cluster_louvain, modularity*) and the detection of hubs (keystone species) was inferred using Kleinberg’s hub centrality score (*hub_score*). To improve visualization, the network was subset according to a minimum node size of 20 (each node is connected by at least 20 edges) and plotted using ggnet2 in R (Villanueva & Chen, 2019).

The top 100 most abundance ASVs (ranked by total abundance, ranging from 15.1-0.1% average relative abundance) along with any suspected human pathogens present at lower abundance (<0.06% relative abundance), representing 105 ASVs in total, were investigated in QTL mapping, along with the leaf surface morphology traits (25 traits), alpha (4) and beta diversity metrics (the first 5 principal components of unweighted UniFrac, weighted UniFrac, Jaccard and Bray-Curtis distances, 20 traits). The method of quantitative trait loci (QTL) mapping was selected according to trait distribution. For normally distributed traits and those which could be normalised by data transformation, mapping was conducted by composite interval mapping in QTL Cartographer v2.5 (North Carolina State University, NC, USA), using the linkage map and genotype data described in Damerum *et al.* (2021). The logarithm of odds (LOD) threshold determining QTL significance at α = 0.05 was determined by permutation with 1,000 iterations and 10 background markers were incorporated into the statistical model as cofactors by forwards stepwise regression with backwards elimination to control for background genetic variation. For traits demonstrating a zero-inflated distribution, which was often observed for ASV counts, a parametric, single-QTL, two-part model was applied using the R package QTL (Broman, 2003, 2009). With this two-part model, individuals with the null phenotype (zero) are considered separately to those with a positive phenotype, which are assumed to follow a normal distribution. Three LOD scores are generated at each genomic position: LODπ; which considers the null phenotype, LODμ; the quantitative phenotype and LODπμ; a sum of both LOD scores and a LODπμ threshold of 3 was used to determine QTL significance. Percentage variance explained was estimated by 1 – 10 ^−2LOD/*n*^, where *n* is the sample size (Broman, 2009). The QTL plot was drawn using circos v0.69-8 (Krzywinski *et al.*, 2009).

### Candidate gene mining

Co-locating QTL, considered interesting for candidate gene mining, were determined by overlap of the 2-LOD intervals (region of the LOD curve in which the LOD score is within 2 of its maximum). The closest marker to this LOD boundary was located and all annotated gene models within this genomic range were retrieved from the *L. sativa* cv. Salinas V8 reference genome (genome ID: 35223) via the CoGe platform (Castillo *et al.*, 2018). cDNA sequences were searched against the *Helianthus annus* genome (HanXRQr1.0) peptide sequences, accessed via Ensembl Plants (http://plants.ensembl.org/index.html) using BLASTx (BLAST version 2.9.0) with an E value cut-off of < 1 × 10^−5^. Significantly enriched gene ontologies were identified using ShinyGO v0.61 (Ge *et al.*, 2020), with an FDR-corrected P-value cut-off of 0.05.

## RESULTS

### Unravelling differences between wild and cultivated lettuce leaf surface morphology and leaf bacterial microbiome

Leaf surface morphology traits for wild and cultivated lettuce parents of the RIL population, differs dramatically (Fig. 1A), such that the cultivated lettuce has larger leaves (t_25_=4.09, P<0.001, Fig. 1B), more circular in shape (t_25_=3.22, P<0.01), with tightly packed, smaller epidermal cells (abaxial: t_20_=−2.68, P<0.05), reduced stomatal index (adaxial: t_21_=−3.46, P<0.01 and abaxial: t_22_=−7.61, P<0.001) and larger stomata (abaxial stomatal length: t_19_=2.22, P−0.05 and adaxial width: t_19_=2.22, P<0.05). Contact angle, used as a measure of leaf surface hydrophobicity, was reduced for the abaxial surface of cultivated lettuce (t_23_=−4.27, P<0.001), suggesting a more hydrophilic leaf surface compared to wild leaves.

**Figure 1.**
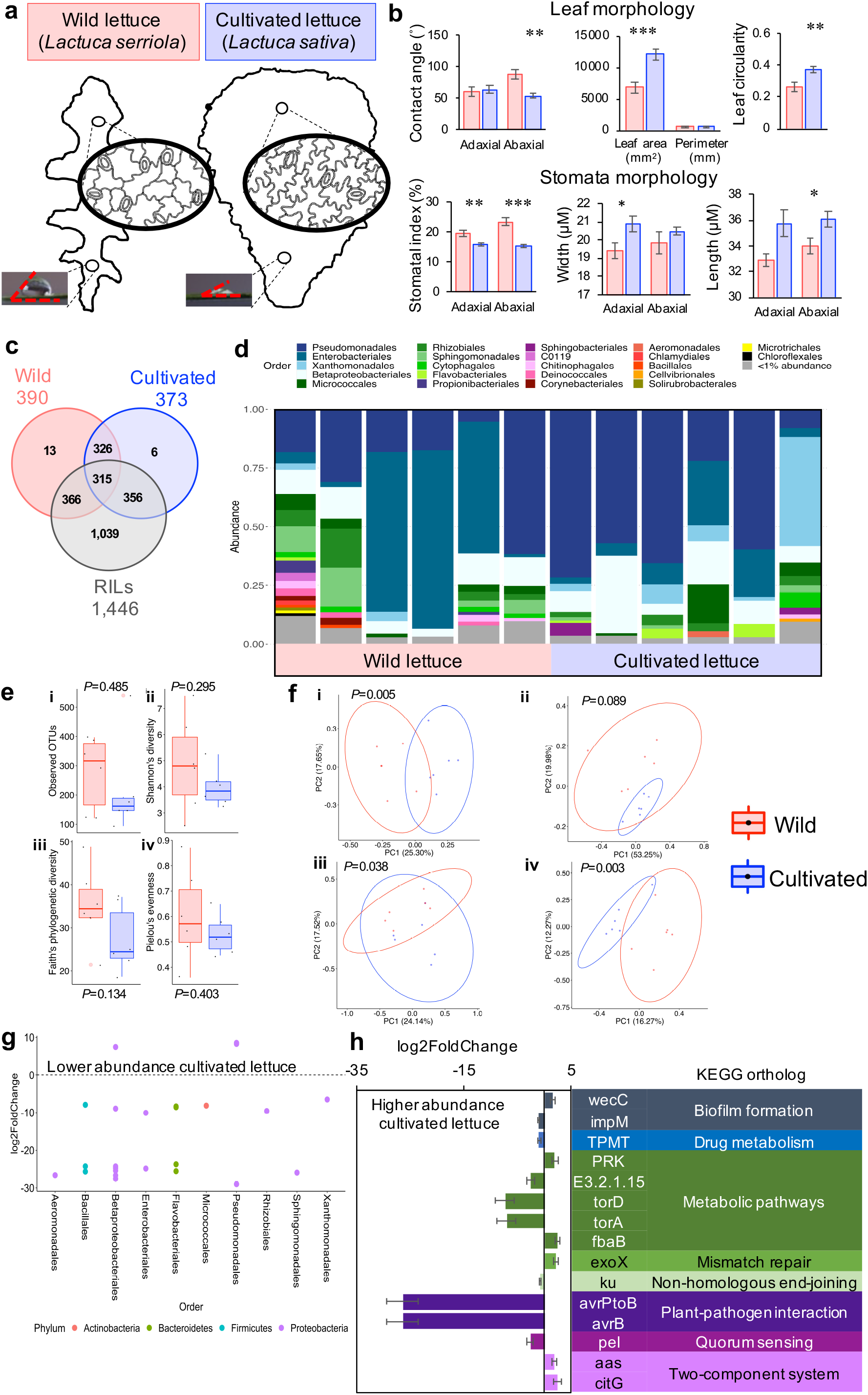
Wild and cultivated lettuce leaf traits and bacterial microbiome. Key leaf surface differences between wild and cultivated lettuce (RIL parents), including leaf size, cell size and stomatal density and leaf surface hydrophobicity (A, B). Venn diagram outlining ASVs present in >5% individuals shared between the wild parent, cultivated parent and RILs (C). Taxa plot demonstrating relative abundance at the Order level (D). Alpha diversity metrics i) observed OTUs, ii) Shannon’s index, iii) Faith’s index and iv) Pielou’s index, with P values indicating results of two-sample t-tests (E). Principal coordinate analysis of beta diversity metrics i) unweighted Unifrac, ii) weighted UniFrac, iii) Bray-Curtis and iv) Jaccard distance, with PERMANOVA P values (F). Differentially abundant ASVs, where each dot represents an ASV coloured according to phylum, with log fold change expressed as wild lettuce compared to cultivated (G). Differentially abundant KEGG orthologs (H) assigned using PICRUSt2 (Douglas *et al.*, 2020), identified using DESeq2 (Love *et al.*, 2014), with bars representing log fold change with standard error, expressed as wild lettuce compared to cultivated.

The leaf bacterial microbiome differed between wild and cultivated lettuce grown in the same environment. 16S profiling identified 437 taxa (amplicon sequence variants, ASVs) present in >5% of the samples, binned at 99% similarity cut-off using the SILVA 132 database, representing 134 uniquely defined genus in 14 phyla (Table S1). Of the 437 ASVs, 326 were shared amongst the parents and 64 and 47 were unique to either wild or cultivated lettuce, respectively (Fig. 1C). The phyllosphere was dominated by proteobacteria (80.7% and 81.1% relative abundance, wild and cultivated lettuce, respectively), largely by the genus *Pseudomonas* (22.5% and 38.9%, wild and cultivated lettuce, respectively) and *Buchnera* (30.3% and 8.7%, wild and cultivated lettuce, respectively), though no significant differences were observed due to high inter-sample variation (Fig. 1D). Actinobacteria was the next most abundant phyla (4.9% and 4.7% relative abundance, wild and cultivated lettuce, respectively), followed by Bacteroidetes (2.4% and 4.4%, wild and cultivated lettuce, respectively). A trend for reduced α diversity of the cultivated parent was observed (Fig. 1E) which was more pronounced when phylogenetic differences were accounted for (Fig. 1Eiii), though not significant. The microbiomes of cultivated and wild lettuce were significantly different when assessed by β diversity measures, including principal coordinate analysis (PCoA) on unweighted UniFrac, Bray-Curtis and Jaccard distances, but not for weighted UniFrac (Fig. 1F), indicating that rare taxa are important in driving the differences in bacterial community structure.

DESeq2 was used to identify differentially abundant ASVs between cultivated and wild lettuce which may be driving these differences in diversity, detecting 25 differentially abundant ASVs in the phyla Proteobacteria (16 features), Bacteroidetes (5), Firmicutes (3) and Actinobacteria (1) (Fig. 1G, Table S2). The majority of these ASVs (22/25) demonstrated increased abundance in cultivated lettuce. To investigate how differences in community composition may translate into differences in community function, functional categories were inferred from taxonomic composition of wild and cultivated lettuce using PICRUSt according to the KEGG Orthology (KO) database. A total of 35 out of 6,477 KO identifiers were differentially abundant between wild and cultivated lettuce using DESeq2 (Fig. 1H, Table S3). Amongst the differentially abundant microbial functions included two KO identifiers (K13452 and K13455) in the plant-pathogen interaction pathway encoding avirulence proteins and another (K11890) in the pathway biofilm formation – *Pseudomonas aeruginosa*, which demonstrated increased abundance in the cultivated lettuce leaf microbiome.

### Characterizing the core leaf bacterial microbiome across a lettuce population

We quantified the leaf microbiome for a diverse population of lettuce RILs composed of 171 individuals assessed in triplicate (n=513) that passed quality control. Quality filtered and rarefied sequences were classified into 9,493 ASVs, binned at 99% similarity cut-off using the SILVA 132 database. A total of 1,084 ASVs could not be assigned a taxonomy, were classified to the kingdom level only or were classified in the kingdom Eukaryota and were searched via BLASTn against the NCBI database. Approximately ~93% of these unassigned sequences had a hit to the *Lactuca sativa* genome, ~5% had no significant similarity to any other sequence and ~2% matched a bacterial hit with low identity and thus were removed from analyses. There were 6,963 ASVs present in <5% of the RILs which were excluded from analyses, resulting in 1,446 ASVs in 18 different phyla and 270 genera.

The core microbiome, defined as the taxa present in >95 % of the leaf samples, was comprised of 45 ASVs belonging to five phyla (Proteobacteria, Actinobacteria, Firmicutes, Chloroflexi and Deinococcus-Thermus), 19 families and 23 genera. Most of these core taxa (32/45) were present at <1% relative abundance. Similar to the RIL parents, the population microbiome was largely represented by Proteobacteria (average relative abundance 80.8%), with Actinobacteria being the next most abundance phylum at ~5.4%, followed by Bacteroidetes (~3.7%), Firmicutes (~1.4%), Deinococcus-Thermus (~1.0%), Chloroflexi (0.8 %) and Acidobacteria (0.6%), though significant between sample variation at the order level was observed (Fig. 2A). When merging ASVs into genus, the most abundant (≥1% relative abundance) were mostly Proteobacteria, including *Pseudomonas* (~36.10%), *Buchnera* (~18.42%), *Alkanindiges* (~4.64%), *Massilia* (3.69%), *Acinetobacter* (3.60%), *Sphingomonas* (3.50%), *Duganella* (2.65%), *Methylobacterium* (2.58%), *Pantoea* (1.89%), *Xanthomonas* (1.58%), *Aquabacterium* (1.35%), *Methylophillus* (1.32%), *Deinococcus* (1.20%), *Chryseobacterium* (1.06%) and *Aquitalea* (1.00%). On average, approximately ~25% of taxa present were low abundance features (<0.1%). Of the 1,446 ASVs, 315 were shared amongst both wild and cultivated parents and 1,039 were unique to the RILs (Fig. 1C). The RILs had increased α diversity compared to the parents (Fig. 2B). Despite similarities in microbiome diversity, variation in ASV abundance was observed across the population (Fig. S1C). Taxa in genera known to be pathogenic to humans were identified, observed in a limited number of samples (5-15%) and at low relative abundance, including a sequence classified as *Escherichia-Shigella* found on leaves of 20/171 RILs at an average of 0.53% relative abundance, two *Clostridium* spp. (13/171 RILs, 0.75% average abundance and 10/171, 0.04% average abundance), *Campylobacter* (10/171 RILs, 0.18% average abundance) and Staphylococcus (27/171 RILs, 0.09% average abundance).

**Figure 2.**
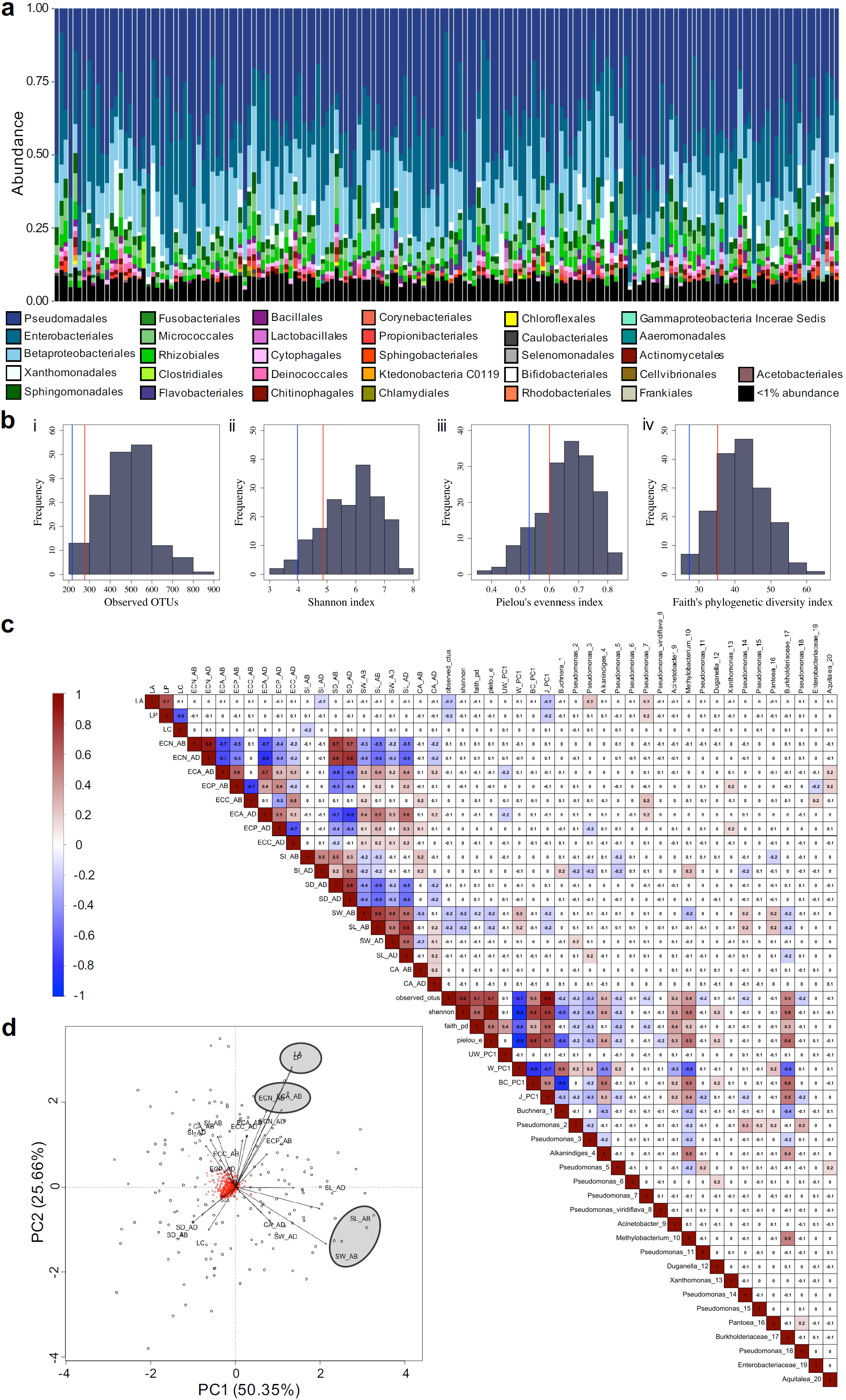
Links between lettuce leaf surface traits and the bacterial microbiome. Taxonomic summary of relative abundance at the order level for the RIL population, where each stack bar represents an individual RIL in the population (n=171; A). Frequency distributions of four α diversity metrics i) observed OTUs, ii) Shannon’s index, iii) Faith’s index and iv) Pielou’s index, with solid horizontal line demonstrating mean trait values for wild (red) and cultivated (blue) lettuce (B). Spearman’s Rank correlation matrix for leaf traits, α diversity metrics, the first principal component of four β diversity metrics and the top 20 most abundant ASVs (C). Coloured boxes indicate significant correlations (P<0.05), with numbers inside denoting the correlation coefficient. Principal component analyses of the ASVs present in >5% of the RILs, with leaf traits significantly correlated with community structure circled in blue (see text for details; D).

**Figure 3.**
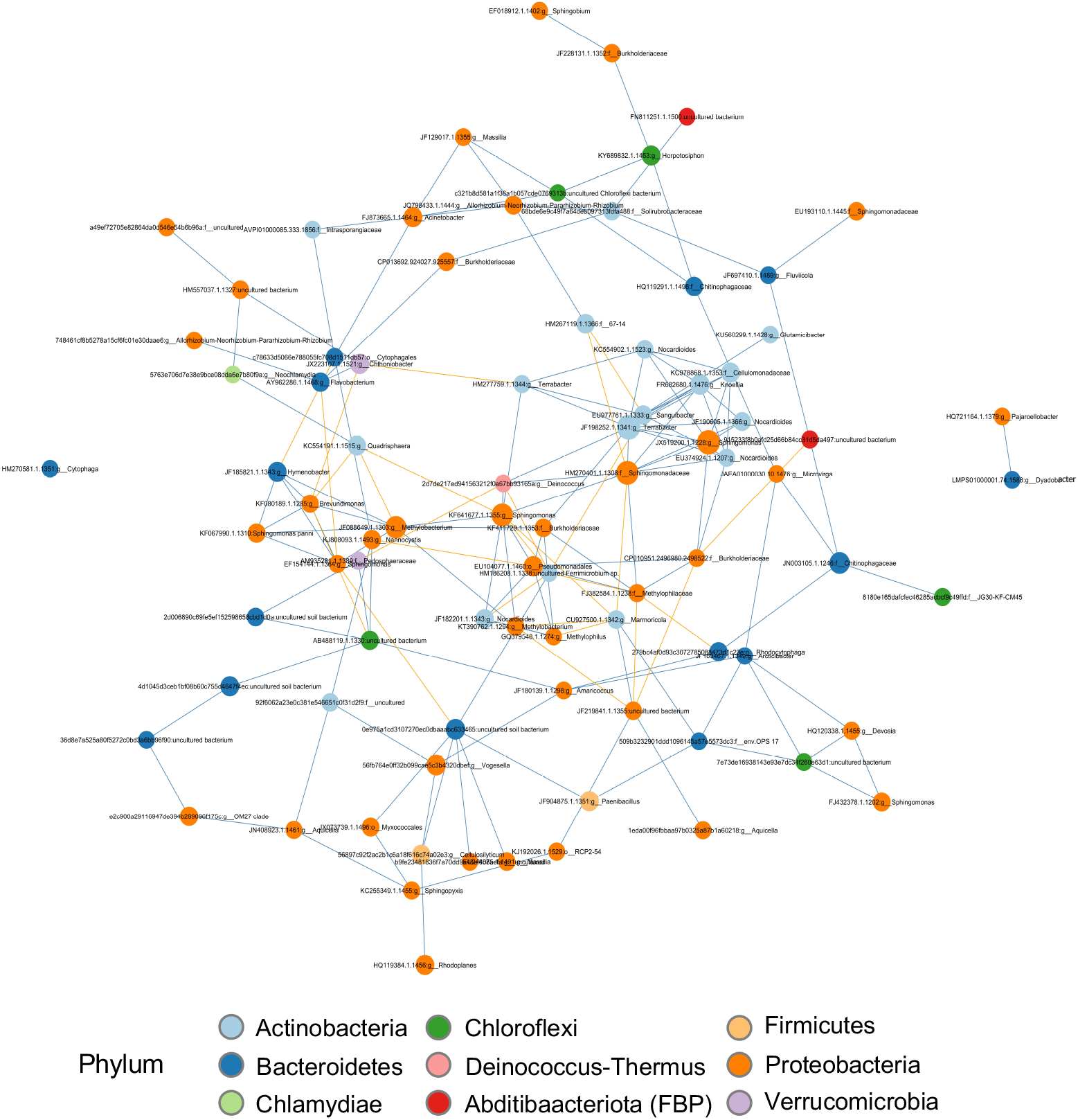
Investigating bacterial cooccurrence networks of the lettuce leaf surface. Nodes of the network represent an ASV, coloured and sized according to phylum and node degree, respectively. The network has been subset to show nodes with a minimum of 20 edges, coloured in blue for positive and yellow for negative edges.

### Leaf surface traits and the bacterial microbiome

Phenotypic variation in the lettuce progeny was observed alongside transgressive segregation above and below parent trait values (Fig. S1), as has been seen for these traits in a previous study (Zhang *et al.*, 2007). Relative to the adaxial (AD) surface, the abaxial (AB) leaf surface had a reduced epidermal cell number (ECN; t_410_=−2.69, *p*<0.01), of similar area (ECA) but a more irregular shape according to epidermal cell circularity (ECC; t_350_=−16.77, *p*<0.001), an increased stomatal density (SD; t_410_=2.97, *p*<0.01) and index (SI; t_382_=9.52, *p*<0.001), but comparable stomata size and a more hydrophobic surface, as measured by contact angle (CA; t_364_=2.60, *p*<0.01). For both the AB and AD surfaces, SD was negatively correlated with stomata size (stomata length; SL and width; SW) and ECA and negatively correlated with ECN (Fig. 2C).

Contrasting leaf surface characteristics across the population were associated with differences in bacterial community composition. Stomata size, but not SD or SI, influenced leaf bacterial α diversity (Fig. 2C). AB SL and SW were negatively correlated with the number of observed OTUs, Shannon diversity and Pielou’s evenness indexes, but no correlation was found with the AD leaf surface stomatal morphology traits. Permutational multivariate analysis of variance (PERMANOVA) of β diversity distances was used to identify leaf traits correlating with β diversity profiles (Table S4). Leaf area (LA) and AB SW and SL varied significantly with all β diversity distances measured (unweighted and weighted UniFrac, Bray-Curtis and Jaccard), explaining between 0.8-2.8% of the total variance (Table S4). Weighted UniFrac distance was significantly influenced by AD ECA (R^2^=0.017, P<0.05), ECN (R^2^=0.029, P<0.01), SD (R^2^=0.016, P<0.05), SL (R^2^=0.021, P<0.01) and SW (R^2^=0.019, P<0.05). AD SI explained 1.4% and 1.1% of the variance in Bray-Curtis and Jaccard distances, respectively (P<0.001; Table S4).

Observing correlations amongst leaf traits and the top 20 most abundant ASVs, LA was positively correlated with two *Pseudomonas* spp. and AD SI was positively correlated with *Buchnera* and *Methylobacterium* spp. and negatively correlated with four *Pseudomonas* spp. (P<0.05; Fig. 2C, Table S5). AB SW was negatively correlated with a *Methylobacterium* spp. and both AB SW and SL were positively correlated with a *Pseudomonas* and *Pantoea* spp. and negatively correlated with a species in the Burkholderiaceae family (P<0.05; Fig. 2C, Table S5). AD SW and SL were each positively correlated with a *Pseudomonas* spp..

Performing principal component analysis (PCA) with the 1,446 ASVs present in >5% of the RILs (Table S6) and applying the *envfit* function (vegan package, R) identified AB SL (R^2^=0.056, P<0.01) and SW (R^2^=0.084, P<0.001), AB ECA (R^2^=0.060, P<0.01) and ECN (R^2^=0.049, P<0.05) and leaf area and perimeter (R^2^=0.129, P<0.001 and R^2^=0.123, P<0.001, respectively) to show significant associations with bacterial community structure (Fig. 2D)

RILs with extreme trait values were further investigated for differentially abundant ASVs and microbial functions using DESeq2 (Love *et al.*, 2014). RILs differing in AB CA, stomatal density, width, and length showed differential abundance of *Pseudomonas*, *Sphingomonas*, *Massillia* and *Pantoea* spp. (Table 1). *Pseudomonas* spp. were less abundant on leaves with reduced AB and AD SD (Table 1; P<0.05 and P<0.001, respectively). The same *Pseudomonas* spp. that had reduced abundance on leaves with low AD SD was found in greater abundance on leaves with reduced AD SL (P<0.001), whilst another *Pseudomonas* spp. was present at lower abundance (P<0.001). Leaves with a low AB SW had a reduction in two *Pseudomonas* spp. (P<0.01) but an increase in one *Pseudomonas* spp. (P<0.001) compared to leaves with increased SW, indicating that different *Pseudomonas* spp. occupy different niches on the leaf surface. Leaves with a lower AB contact angle (CA), indicating a more hydrophilic leaf surface, had a reduced abundance of *Sphingomonas* spp. (P<0.01).

**Table 1.**
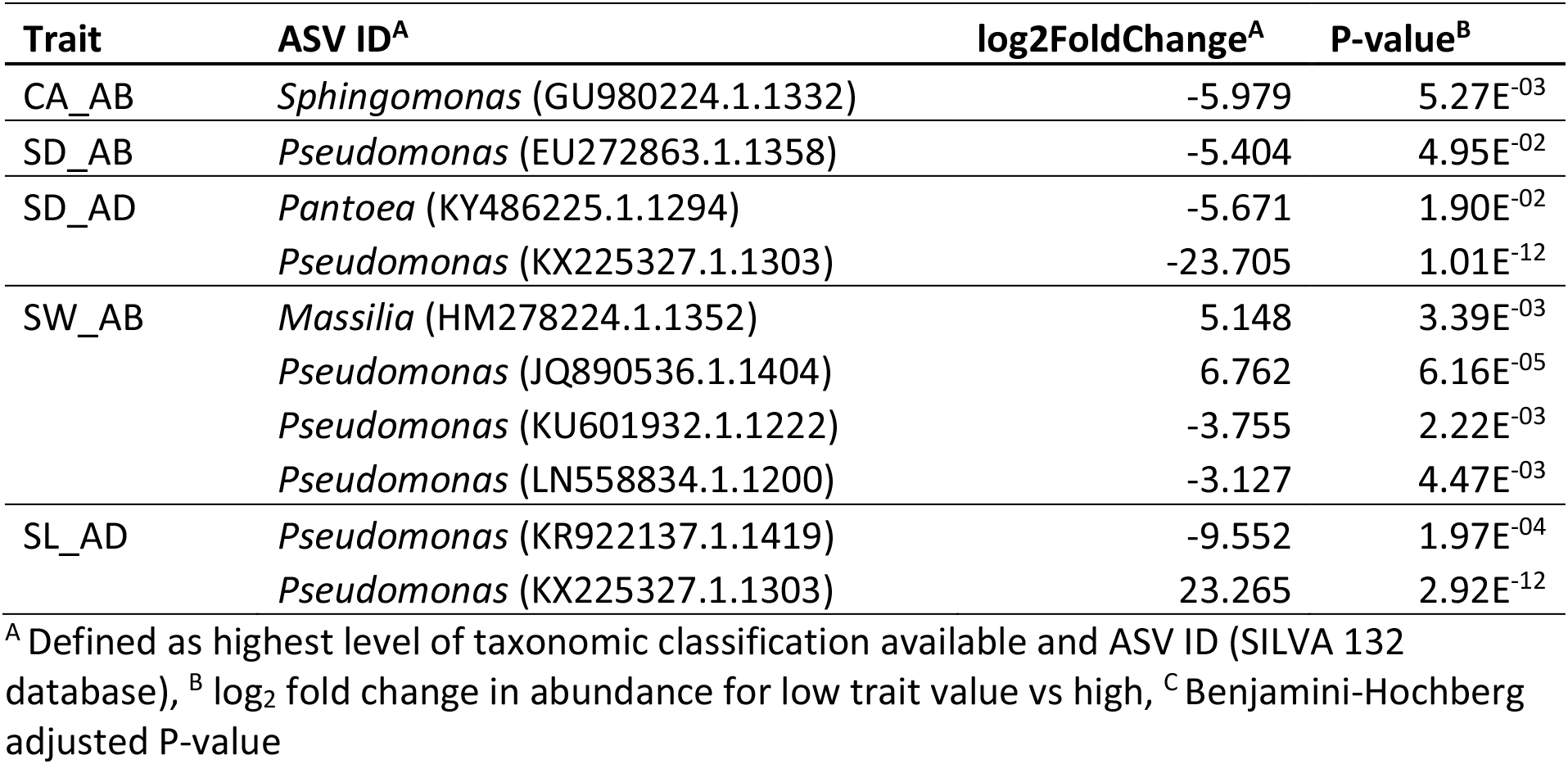
ASVs differentially abundant between extreme RILs with low vs high trait values

Comparing the predictive functional profiles of microbial communities determined using PICRUSt2 between leaves with extreme trait values identified significantly abundant KEGG Orthologs (KOs) for abaxial surface traits (CA, SD, SI, SL and SW) and whole LA (Table S8). 2,391 KOs were differentially abundant between leaves with low vs high AB SW and 1,749 for low vs high LA, classified most frequently into pathways including secondary metabolite biosynthesis (ko01100), microbial metabolism in diverse environments (ko01110), ABC transporters (ko02010), two-component system (ko02020), quorum sensing (ko02024), bacterial secretion system (ko03070) and biofilm formation – *Pseudomonas aeruginosa* (ko02025). Two KOs, a molecular chaperone (HSP90A) and an outer membrane protein (tolC; K04079 and K12340, respectively) in the plant-pathogen interaction pathway were found at reduced abundance on leaves with lower AB SW (Table S8, P<0.05). Seventeen genes involved in chemotaxis were differentially abundant in leaves with smaller stomata width, generally observed at reduced abundance (13/17 genes) and all six genes involved in chemotaxis in leaves with a smaller area were observed at a reduced abundance. Numerous genes involved in drug resistance or transportation were differentially abundant in leaves with lower AB SW and LA, with 15/35 and 9/20, respectively, demonstrating reduced abundance at lower trait values. Genes functioning in quorum sensing (K07782, K13651, K19734) were observed in lower abundance on leaves with lower AB SW and those functioning in biofilm formation where more abundant on bigger leaves (K19449, K13654, K13650).

### Bacterial species influencing the leaf surface community composition

The most abundant ASV, in the genus *Buchnera* and multiple ASVs classified as *Pseudomonas* spp. were negatively correlated with all α diversity measures, whereas ASVs in the genera *Alkanindiges*, *Acinetobacter*, *Aquabacterium*, *Duganella*, *Methylobacterium* a, *Methylophils* and the family Burkholderiaceae were positively correlated with α diversity (Fig. 2C, Table S5). *Pseudomonas* spp. were generally positively correlated with other *Pseudomonas* spp. and negatively correlated with *Alkanindiges* spp. and *Methylobacterium* spp..

To investigate bacterial dynamics more rigorously, co-occurrences amongst the lettuce leaf bacterial community were explored in network analyses using SPIEC-EASI (Kurtz *et al.*, 2015), which utilises a centered log-ratio transformation appropriate for compositional data. From the 1,446 nodes representing the ASVs present in >5% RILs, a total of 8,024 edges were detected, the majority of which were positive (7,108), indicating a co-presence and 916 were negative (co-occurrence observed less frequently than expected by random). The average node degree (number of connections each node has) was 11.10 and the average path length (number of edges between nodes) was 3.38. The Louvain algorithm was implemented to identify 16 communities (Table S9) with a modularity of 0.36 and a clustering coefficient of 0.05, indicating the presence of numerous, weak ties between nodes. Taxa from the same genera generally did not demonstrate shared community membership, suggesting the microbiome structure is more influenced by biological functions than phylogeny, which has previously been observed (Copeland *et al.*, 2015).

Hub scores were computed to identify keystone taxa which have the largest influence on overall bacterial community structure (Table S9). As previously observed (Roman-Reyna *et al.*, 2019), hub score did not correlate with taxa abundance, with the taxa demonstrating hub scores >0.7 contributing to <0.5% of the relative abundance. A *Terrabacter* spp. in the phylum Actinobacteria, had the highest hub score of 1, followed by taxa in the phylum proteobacteria including an ASV in the Sphingomonadaceae family (hub score = 0.90) and two *Sphingomonas* spp. (0.80 and 0.78). These keystone taxa demonstrated more positive interactions (21-35) than negative (7-14). Despite the high abundance of *Pseudomonas* spp. average hub score was <0.05 and average node degree was <5, suggesting low connectivity with other species. The most abundant taxa, identified to negatively correlate with diversity metrics, demonstrated more positive correlation with taxa than negative (Table S9). *Buchnera* spp. was negatively correlated with a taxon in the Burkholderiaceae family (1.2% abundance), *Terrabacter* (0.1% abundance), *Nocardiooides* (0.07% abundance), *Sphingomonas* (0.05% abundance) and positively correlated with 8 ASVs low abundance ASVs (<0.02%), including *Sphingomonas* spp. and taxa in the phylum Bacteroidetes (*Dyadobacter*, *Mucilaginibacter*, and two taxa in the Chitinophagaceae family; Table S9). The most abundant *Pseudomonas* spp. demonstrated no negative correlations and was positively correlated with two other Pseudomonas spp. (3.1% and <0.01% abundance).

### Genetic architecture of the lettuce bacterial microbiome

Quantitative trait loci (QTL) mapping was conducted to identify lettuce genomic loci affecting leaf surface traits and bacterial microbiome composition. A total of 25 leaf surface traits, 24 alpha and beta diversity metrics and 105 selected ASVs were investigated in QTL analyses. Collectively, 306 QTL were identified across the nine linkage groups (LGs) for leaf surface traits (81 QTL), diversity metrics (92) and for ASVs (133) explaining between 2.5-37.4% of the phenotypic variance (Fig. 4, Table S10).

**Figure 4.**
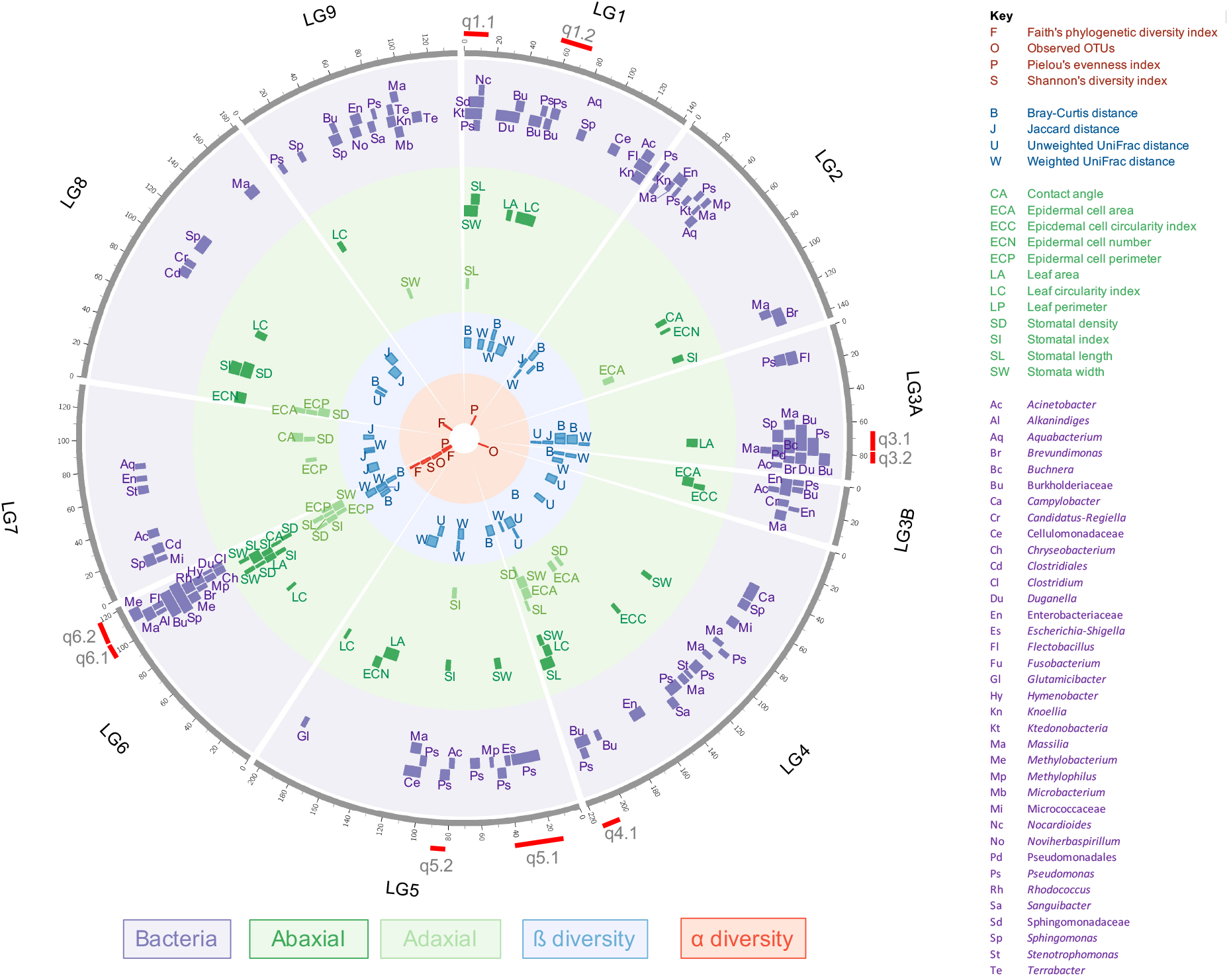
Quantitative trait loci mapping of the lettuce bacterial phyllosphere microbiome. Distribution of QTL for bacteria (purple), abaxial and adaxial leaf traits (green) and alpha (red) and beta (blue) diversity metrics. Boxes represent QTL 1-LOD interval. Outer grey circle represents linkage group position in the lettuce genome in cM. Red dashes indicate regions considered for candidate gene mining. QTL details are given in Table 2 and Table S10.

QTL for leaf surface traits often demonstrated clustering and were observed to co-locate for both the AB and AD surfaces for stomatal traits on LGs 1, 4, 5 and 6. QTL for stomata traits and leaf size and shape frequently mapped to the same genomic position as those for α and β diversity metrics (Fig. 4, Table 2; q1.1-2, q3.1, q4.1, q5.2 and q6.1-2). In a region on LG 1 (q1.1), QTL for SL (AB and AD) and SW (AB) mapped to the same position as QTL for taxa in the genera *Pseudomonas*, *Nocardioides*, the Sphingomonadaceae family (identified as a keystone taxa in co-occurrence network analyses) and in the class Ktedonobacteria. A region on LG 3.1 (q3.1) harboured QTL for leaf area mapping to the same position as *Buchnera*, *Pseudomonas*, *Sphingomonas*, two taxa in the family Burkholderiaceae and β diversity metrics unweighted UniFrac and Bray-Curtis. In another region (q3.2) leaf perimeter mapped to the same position as QTL for a *Pseudomonas* spp. and Bray-Curtis distance. Two regions on LG 6 from 98.4-103.8 cM (q6.1) and 107.2-117.7 (q6.2) contained overlapping QTL for AB and AD stomatal traits, multiple α and β diversity indices and several genera of bacteria, including a *Sphingomonas* spp. identified as a keystone taxa and *Methylobacterium* (q6.1) and *Alkanindiges*, *Brevundimonas*, *Chryseobacterium*, *Duganella*, *Hymenobacter, Flectobacillus, Masilia, Methylophilius, Rhodococcus* and *Clostridium* (q6.2; Fig. 4, Table 2). QTL for keystone taxa *Terrabacter* spp. (LG 9), Sphingomonadaceae (LG 1) and *Sphingomonas* spp. (LGs 6 and 8) were identified. The individual taxa with the most QTL detected was a member of the Burkholderiaceae family, with seven QTL observed on LGs 1, 3,4,6 and 9. QTL for *Pseudomonas* spp., the most abundant genus, co-located with QTL for stomata size, density or index on LGs 1 (q1.1), 4 (q4.1) and 5 (q5.1-2) and leaf size and shape on LGs 1 (q1.2), 3 (q3.1-2) and 4 (q4.1). Despite low abundance and occurrence, QTL for potential pathogenic bacteria were identified on LGs 4 (*Campylobacter* spp.), 5 (*Escherichia-Shigella* spp.; q5.1) and 6 (*Clostridium* spp.; q6.2). A QTL for AB SW co-located with a QTL for *Escherichia-Shigella* spp. and *Pseudomonas* spp. on LG 5 (q5.1).

**Table 2.** QTL details for investigated candidate regions. Extended table given in Table S10.

| Region | Trait<sup>A</sup> | Fig. 4 ID | LG | Position (cM) | Position (Mbp) | LOD<sup>B</sup> | R<sup>2</sup> (%)<sup>C</sup> | LOD2 lower (cM)<sup>D</sup> | LOD2 upper (cM)<sup>D</sup> | QTL model<sup>E</sup> |
| --- | --- | --- | --- | --- | --- | --- | --- | --- | --- | --- |
| q1.1 | JF135626.1.1327:c__Ktedonobacteria | Kt | 1 | 3.6 | 4.2 | 3.5 | 7.7 | 0.3 | 9.3 | CIM |
| q1.1 | Stomatal width (AB) | SW | 1 | 6.4 | 7.5 | 4.4 | 4.9 | 3.7 | 9.3 | CIM |
| q1.1 | LN558834.1.1200:g__Pseudomonas | Ps | 1 | 9.3 | 9.8 | 3.5 | 6.3 | 6.4 | 16.8 | CIM |
| q1.1 | Bray-Curtis distance (PCoA 3) | B | 1 | 10.3 | 9.8 | 4.1 | 7.4 | 0.0 | 15.1 | CIM |
| q1.1 | JF135626.1.1327:c__Ktedonobacteria | Kt | 1 | 11.3 | 9.8 | 3.6 | 7.4 | 9.3 | 13.3 | CIM |
| q1.1 | HM270401.1.1308:f__Sphingomonadaceae | Sd | 1 | 11.4 | 11.8 | 3.7 | 7.1 | 0.3 | 14.3 | CIM |
| q1.1 | EU374924.1.1207:g__Nocardioides | Nc | 1 | 11.4 | 11.8 | 4.2 | 7.5 | 5.3 | 14.4 | CIM |
| q1.1 | Stomatal length (AD) | SL | 1 | 11.4 | 11.8 | 3.5 | 4.6 | 3.1 | 15.3 | CIM |
| q1.1 | Stomatal width (AB) | SW | 1 | 11.9 | 13.4 | 5.5 | 5.9 | 9.3 | 15.1 | CIM |
| q1.1 | Stomatal length (AB) | SL | 1 | 11.9 | 13.4 | 5.2 | 6.5 | 4.1 | 15.6 | CIM |
| q1.2 | HF572845.1.1261:g__Pseudomonas | Ps | 1 | 55.2 | 56.2 | 4.7 | 11.8 | 44.2 | 64.9 | ZIM |
| q1.2 | HQ018861.1.1234:g__Pseudomonas | Ps | 1 | 55.3 | 56.2 | 4.5 | 7.8 | 54.2 | 56.1 | CIM |
| q1.2 | Leaf circularity index | LC | 1 | 55.4 | 59.1 | 15.6 | 17.0 | 53.5 | 88.5 | CIM |
| q3.1 | KF441583.1.1479:g__Sphingomonas | Sp | 3.1 | 71.2 | 96.3 | 3.4 | 5.3 | 60.7 | 78.1 | CIM |
| q3.1 | GU086441.1.1495:f__Burkholderiaceae | Bu | 3.1 | 71.2 | 96.3 | 4.7 | 9.0 | 69.7 | 77.8 | CIM |
| q3.1 | Unweighted UniFrac distance (PCoA 4) | U | 3.1 | 71.2 | 96.3 | 2.5 | 2.9 | 70.8 | 77.8 | CIM |
| q3.1 | Bray-Curtis distance (PCoA 1) | B | 3.1 | 71.2 | 96.3 | 3.7 | 6.9 | 63.0 | 77.8 | CIM |
| q3.1 | KM187229.1.1392:f__Burkholderiaceae | Bu | 3.1 | 71.2 | 96.3 | 3.6 | 9.2 | 43.0 | 80.0 | ZIM |
| q3.1 | KP295476.1.1410:g__Pseudomonas | Ps | 3.1 | 72.2 | 96.3 | 4.2 | 7.2 | 69.7 | 77.8 | CIM |
| q3.1 | Leaf area | LA | 3.1 | 72.2 | 96.3 | 9.4 | 12.2 | 71.2 | 77.5 | CIM |
| q3.1 | JX998102.1.1391:g__Buchnera | Bc | 3.1 | 72.5 | 97.4 | 3.2 | 5.6 | 64.1 | 82.3 | CIM |
| q3.1 | Bray-Curtis distance (PCoA 5) | B | 3.1 | 74.5 | 97.4 | 5.7 | 10.2 | 69.7 | 77.8 | CIM |
| q3.2 | Bray-Curtis distance (PCoA 1) | B | 3.1 | 78.2 | 102.9 | 3.4 | 6.3 | 77.8 | 81.3 | CIM |
| q3.2 | Leaf perimeter | LP | 3.1 | 79.2 | 102.9 | 9.6 | 13.9 | 74.3 | 79.9 | CIM |
| q3.2 | KP295476.1.1410:g__Pseudomonas | Ps | 3.1 | 79.9 | 107.0 | 4.3 | 7.2 | 77.8 | 81.7 | CIM |
| q3.2 | EU104077.1.1460:o__Pseudomonadales | Pd | 3.1 | 79.9 | 107.0 | 3.2 | 6.0 | 72.5 | 81.0 | CIM |
| q3.2 | Bray-Curtis distance (PCoA 5) | B | 3.1 | 79.9 | 107.0 | 7.2 | 12.0 | 77.8 | 82.1 | CIM |
| q3.2 | KF101691.1.1351:g__Massilia | Ma | 3.1 | 81.0 | 111.0 | 4.6 | 8.0 | 79.3 | 84.0 | CIM |
| q3.2 | GU086441.1.1495:f__Burkholderiaceae | Bu | 3.1 | 81.0 | 111.0 | 6.7 | 12.2 | 79.4 | 85.0 | CIM |
| q3.2 | KM021214.1.1333:g__Duganella | Du | 3.1 | 81.7 | 113.8 | 4.7 | 8.8 | 78.6 | 84.0 | CIM |
| q3.2 | Weighted UniFrac distance (PCoA 2) | W | 3.1 | 81.7 | 113.8 | 9.1 | 13.9 | 77.9 | 84.0 | CIM |
| q3.2 | Jaccard distance (PCoA 2) | J | 3.1 | 81.7 | 113.8 | 7.9 | 13.3 | 79.3 | 82.5 | CIM |
| q4.1 | Leaf circularity index | LC | 4 | 209.2 | 356.0 | 5.5 | 5.3 | 203.8 | 211.9 | CIM |
| q4.1 | Epidermal cell number (AD) | ECN | 4 | 210.6 | 359.7 | 4.1 | 5.7 | 206.7 | 214.6 | CIM |
| q4.1 | Stomatal density (AD) | SD | 4 | 210.6 | 359.7 | 3.2 | 4.5 | 205.3 | 214.3 | CIM |
| q4.1 | Stomatal width (AB) | SW | 4 | 210.6 | 359.7 | 14.2 | 17.4 | 207.8 | 216.4 | CIM |
| q4.1 | Stomatal length (AB) | SL | 4 | 210.6 | 359.7 | 8.7 | 11.6 | 206.1 | 213.4 | CIM |
| q4.1 | Stomatal length (AD) | SL | 4 | 210.6 | 359.7 | 8.2 | 11.9 | 209.6 | 213.2 | CIM |
| q4.1 | JQ890536.1.1404:g__Pseudomonas | Ps | 4 | 210.8 | 359.7 | 3.8 | 9.8 | 200.6 | 216.3 | ZIM |
| q4.1 | KF411729.1.1353:f__Burkholderiaceae | Bu | 4 | 211.7 | 361.6 | 3.7 | 6.3 | 206.2 | 217.8 | CIM |
| q4.1 | Stomatal width (AD) | SW | 4 | 211.9 | 366.0 | 11.2 | 16.4 | 207.1 | 213.9 | CIM |
| q4.1 | Weighted UniFrac distance (PCoA 4) | W | 4 | 211.9 | 366.0 | 3.2 | 5.6 | 205.4 | 218.4 | CIM |
| q5.1 | EU434431.1.1200:g__Pseudomonas | Ps | 5 | 25.5 | 38.0 | 3.0 | 7.8 | 0.5 | 202.6 | ZIM |
| q5.1 | Stomatal width (AB) | SW | 5 | 36.3 | 52.3 | 3.6 | 4.3 | 33.3 | 39.9 | CIM |
| q5.1 | JF712661.1.1208:g__Escherichia-Shigella | Es | 5 | 39.5 | 60.7 | 4.2 | 10.7 | 35.5 | 44.9 | ZIM |
| q5.2 | Weighted UniFrac distance (PCoA 3) | W | 5 | 83.0 | 158.8 | 9.1 | 15.4 | 81.3 | 84.7 | CIM |
| q5.2 | Weighted UniFrac distance (PCoA 5) | W | 5 | 83.0 | 158.8 | 11.8 | 16.2 | 81.7 | 85.5 | CIM |
| q5.2 | JN406504.1.1364:Pseudomonas viridiflava | Ps | 5 | 85.4 | 160.4 | 6.1 | 10.9 | 82.7 | 87.7 | CIM |
| q5.2 | Stomatal index (AD) | SI | 5 | 85.4 | 160.4 | 4.4 | 6.1 | 83.0 | 90.2 | CIM |
| q5.2 | Stomatal index (AB) | SI | 5 | 87.5 | 162.0 | 4.1 | 4.4 | 86.4 | 89.2 | CIM |
| q6.1 | Stomatal index (AB) | SI | 6 | 98.4 | 155.5 | 7.6 | 8.2 | 95.2 | 101.3 | CIM |
| q6.1 | Stomatal index (AD) | SI | 6 | 98.4 | 155.5 | 8.7 | 13.8 | 98.2 | 100.6 | CIM |
| q6.1 | Stomatal density (AB) | SD | 6 | 98.4 | 155.5 | 3.8 | 5.5 | 98.2 | 101.3 | CIM |
| q6.1 | Stomatal density (AD) | SD | 6 | 98.4 | 155.5 | 4.6 | 6.5 | 98.2 | 101.3 | CIM |
| q6.1 | Leaf area | LA | 6 | 98.4 | 155.5 | 3.4 | 3.9 | 97.2 | 103.2 | CIM |
| q6.1 | Stomatal width (AB) | SW | 6 | 98.4 | 155.5 | 4.0 | 5.3 | 98.2 | 101.3 | CIM |
| q6.1 | KF411729.1.1353:f__Burkholderiaceae | Bu | 6 | 101.3 | 157.6 | 3.3 | 5.8 | 100.4 | 104.7 | CIM |
| q6.1 | KF641677.1.1355:g__Sphingomonas | Sp | 6 | 101.3 | 157.6 | 4.0 | 7.9 | 93.9 | 103.2 | CIM |
| q6.1 | Observed OTUs index | O | 6 | 101.3 | 157.6 | 6.8 | 12.3 | 98.8 | 104.3 | CIM |
| q6.1 | Shannon's diversity index | S | 6 | 101.3 | 157.6 | 4.5 | 9.0 | 100.4 | 104.7 | CIM |
| q6.1 | Faith's phylogenetic diversity index | F | 6 | 101.3 | 157.6 | 7.8 | 13.8 | 99.1 | 102.3 | CIM |
| q6.1 | Jaccard distance (PCoA 1) | J | 6 | 101.3 | 157.6 | 3.9 | 7.4 | 97.2 | 106.5 | CIM |
| q6.1 | Weighted UniFrac distance (PCoA 1) | W | 6 | 102.3 | 157.6 | 8.1 | 16.0 | 99.2 | 104.1 | CIM |
| q6.1 | JF088649.1.1303:g__Methylobacterium | Me | 6 | 103.2 | 159.2 | 6.8 | 11.5 | 101.1 | 104.7 | CIM |
| q6.1 | KF441583.1.1479:g__Sphingomonas | Sp | 6 | 103.2 | 159.2 | 4.7 | 8.0 | 101.0 | 104.7 | CIM |
| q6.1 | Epidermal cell perimeter (AD) | ECP | 6 | 103.2 | 159.2 | 4.2 | 6.3 | 97.8 | 104.7 | CIM |
| q6.1 | Bray-Curtis distance (PCoA 1) | B | 6 | 103.2 | 159.2 | 2.5 | 4.6 | 100.5 | 104.7 | CIM |
| q6.1 | Stomatal index (AD) | SI | 6 | 103.8 | 161.3 | 4.9 | 8.1 | 103.2 | 103.9 | CIM |
| q6.1 | Stomatal length (AB) | SL | 6 | 103.8 | 161.3 | 4.7 | 6.2 | 99.9 | 104.7 | CIM |
| q6.2 | KF101569.1.1356:g__Alkanindiges | Al | 6 | 107.2 | 170.0 | 4.2 | 7.3 | 104.7 | 109.4 | CIM |
| q6.2 | KF067990.1.1310:s__Sphingomonas panni | Sp | 6 | 107.2 | 170.0 | 5.5 | 9.1 | 105.2 | 108.6 | CIM |
| q6.2 | JF237686.1.1306:g__Sphingomonas | Sp | 6 | 107.2 | 170.0 | 8.4 | 13.8 | 105.8 | 118.8 | CIM |
| q6.2 | KF080189.1.1285:g__Brevundimonas | Br | 6 | 107.2 | 170.0 | 4.9 | 8.6 | 104.7 | 108.6 | CIM |
| q6.2 | AB696367.1.1332:g__Massilia | Ma | 6 | 107.2 | 170.0 | 6.2 | 10.2 | 105.5 | 109.9 | CIM |
| q6.2 | Observed OTUs index | O | 6 | 107.2 | 170.0 | 4.6 | 8.2 | 106.5 | 110.2 | CIM |
| q6.2 | Bray-Curtis distance (PCoA 3) | B | 6 | 107.2 | 170.0 | 3.9 | 6.5 | 104.7 | 113.5 | CIM |
| q6.2 | Weighted UniFrac distance (PCoA 1) | W | 6 | 108.2 | 170.0 | 8.2 | 15.6 | 106.2 | 111.3 | CIM |
| q6.2 | Contact angle (AB) | CA | 6 | 108.6 | 170.5 | 5.1 | 6.3 | 106.6 | 110.2 | CIM |
| q6.2 | Stomatal length (AB) | SL | 6 | 109.4 | 172.2 | 7.6 | 9.8 | 106.8 | 113.5 | CIM |
| q6.2 | FQ660167.1.1362:g__Methylophilus | Mp | 6 | 110.2 | 174.1 | 4.0 | 7.4 | 108.6 | 118.2 | CIM |
| q6.2 | Stomatal density (AB) | SD | 6 | 110.2 | 174.1 | 6.8 | 9.9 | 108.8 | 112.7 | CIM |
| q6.2 | Stomatal density (AD) | SD | 6 | 110.2 | 174.1 | 4.4 | 6.3 | 106.2 | 111.7 | CIM |
| q6.2 | Stomatal width (AB) | SW | 6 | 110.2 | 174.1 | 12.9 | 15.3 | 108.0 | 112.0 | CIM |
| q6.2 | Stomatal width (AD) | SW | 6 | 110.2 | 174.1 | 7.2 | 10.3 | 109.4 | 112.9 | CIM |
| q6.2 | Stomatal length (AD) | SL | 6 | 110.2 | 174.1 | 7.1 | 10.1 | 107.5 | 111.7 | CIM |
| q6.2 | Weighted UniFrac distance (PCoA 4) | W | 6 | 110.2 | 174.1 | 4.9 | 8.7 | 108.6 | 111.2 | CIM |
| q6.2 | KF441583.1.1479:g__Sphingomonas | Sp | 6 | 111.2 | 174.1 | 7.3 | 12.1 | 108.6 | 114.3 | CIM |
| q6.2 | LN890295.1.1403:g__Flectobacillus | Fl | 6 | 111.2 | 174.1 | 5.9 | 10.7 | 108.6 | 112.9 | CIM |
| q6.2 | Shannon's diversity index | S | 6 | 111.2 | 174.1 | 5.5 | 10.7 | 110.2 | 112.2 | CIM |
| q6.2 | Faith's phylogenetic diversity index | F | 6 | 111.2 | 174.1 | 4.5 | 8.4 | 109.4 | 114.6 | CIM |
| q6.2 | JF088649.1.1303:g__Methylobacterium | Me | 6 | 111.7 | 176.5 | 11.1 | 17.8 | 110.2 | 114.7 | CIM |
| q6.2 | KF411729.1.1353:f__Burkholderiaceae | Bu | 6 | 111.7 | 176.5 | 3.8 | 6.4 | 109.4 | 117.7 | CIM |
| q6.2 | KU515242.1.1490:g__Massilia | Ma | 6 | 111.7 | 176.5 | 4.8 | 8.1 | 108.6 | 112.9 | CIM |
| q6.2 | JF082030.1.1345:g__Chryseobacterium | Ch | 6 | 111.7 | 176.5 | 3.6 | 6.8 | 107.2 | 112.9 | CIM |
| q6.2 | DQ846830.1.1264:g__Rhodococcus | Rh | 6 | 111.7 | 176.5 | 8.9 | 15.9 | 110.3 | 116.9 | CIM |
| q6.2 | JF185821.1.1343:g__Hymenobacter | Hy | 6 | 111.7 | 176.5 | 6.1 | 10.5 | 109.0 | 117.0 | CIM |
| q6.2 | Pielou's evenness index | Ps | 6 | 111.7 | 176.5 | 4.7 | 8.7 | 106.4 | 118.3 | CIM |
| q6.2 | FJ357796.1.1509:g__Clostridium sensu stricto 1 | Cl | 6 | 114.0 | 180.1 | 5.1 | 12.8 | 112.9 | 114.8 | ZIM |
| q6.2 | KU514606.1.1492:g__Duganella | Du | 6 | 114.8 | 180.1 | 5.4 | 10.0 | 112.9 | 117.0 | CIM |
| q6.2 | KT390762.1.1294:g__Methylobacterium | Me | 6 | 114.8 | 180.1 | 7.5 | 14.0 | 109.4 | 117.6 | CIM |
| q6.2 | Weighted UniFrac distance (PCoA 1) | W | 6 | 114.8 | 180.1 | 2.8 | 4.8 | 112.9 | 119.6 | CIM |
| q6.2 | Epidermal cell perimeter (AD) | ECP | 6 | 115.8 | 180.1 | 5.9 | 9.4 | 114.4 | 119.1 | CIM |
| q6.2 | Stomatal width (AD) | SW | 6 | 117.7 | 183.6 | 9.9 | 13.8 | 115.3 | 120.5 | CIM |

<sup>A</sup> Trait ID, comprised of ASV ID (SILVA 132 database) followed by taxonomic rank and taxa ID, <sup>B</sup> logarithm of odds score, <sup>C</sup> percent variance explained by the QTL, <sup>D</sup> lower and upper LOD2 QTL intervals, <sup>E</sup> model used for QTL mapping

Nine genomic regions demonstrating interesting QTL overlap were investigated for underlying candidate genes and significantly enriched gene ontology (GO) terms (Table 2). On average, ~519 putative genes were identified within candidate regions, ~85% of which could be assigned an annotation and ~90% were characterised (Table S11). Commonly enriched GO terms included those involved in cell wall assembly or modification (q1.1-2, q4.1), in response to stimuli and stress (q1.2, q3.1, q5.1 and q6.1), defence response (q1.2 and q6.2), hormone-mediated signalling and biosynthesis (q1.2, q3.1-2, q6.2), the regulation of metabolic processes (q1.1-2, q3.1, q4.1, q5.1 and q6.1) and anatomical structure development (q3.1-2, q4.1, q5.1, q6.2; Table S12). Frequently observed gene candidates include pentatricopeptide repeat family proteins involved in organellar RNA metabolism (Barkan & Small, 2014; q1.1, 1.2, q3.1, 3.2, q4.1, q5.1, q6.1 and q6.2), leucine-rich repeat receptor-like protein kinases with roles in development and stress response (Dufayard *et al.*, 2017; q1.1 and q3.1), glycosyltransferase (q1.1, q1.2, q3.1, q5.1, q6.1 and q6.2), pectinesterase (q1.2, q4.1 and q6.2), cellulose synthase (q1.1, q1.2, q4.1, q5.1, q5.2 and q6.1) and genes involved in ethylene and auxin response (all regions investigated). Leucine-rich repeat domain containing proteins (all regions investigated) and lectin domain-containing proteins were identified (q1.1-2, q3.1, q5.1-2 and q6.1), which have roles as immune receptors (Lannoo & Van Damme, 2014). Other notable candidates include a defensin-like protein which have antimicrobial activity (Sathoff *et al.*, 2020; q5.2), a putative *TRICHOME-BIREFRINGENCE-LIKE* gene involved in the deposition of callose and an epidermal patterning factor-like gene, which influence the development of stomata in the upper epidermis (q6.2).

## DISCUSSION

The leaf surface provides the first physical barrier to aerial microbes and is crucial in modulating microbial community structure, through the control of plant-microbe and microbe-microbe interactions. Here we investigated the relationship between host plant and bacterial microbiome genetics and have identified together, for the first time, the genetic architecture of these leaf and microbial traits that may be of wide-ranging significance for understanding the lettuce holobiont and in identifying future targets for lettuce breeding of engineered ‘good’ leaf microbiomes.

Our data suggest that domestication has drastically altered lettuce leaf surface characteristics with cultivated lettuce, characterised by larger leaves (representing yield) and fewer larger stomata. A reduction in α diversity in the cultivated lettuce leaf microbiome was observed, but this finding is in contrast with previous observations of the lettuce bacterial rhizosphere microbiome (Cardinale *et al.*, 2015), where an increased number of taxa were observed in the cultivated lettuce. How domestication of food crops has impacted the associated microbiomes still remains to be fully elucidated and it could be as for humans, where genome-wide association studies (GWAS) have identified environmental factors to collectively have a larger effect on microbiome composition than host genetics (Beilsmith *et al.*, 2019), this also holds for plant genotypes. Differences in β diversity were observed for qualitative metrics, which measure shared features (Jaccard distance) and also account for phylogenetic diversity (unweighted UniFrac), but do not consider taxa abundance, and for quantitative measures which account for abundances (Bray-Curtis), but not when phylogeny was also considered (Weighted UniFrac). This indicates that low abundance features are important in driving the differences in bacterial community structure between wild and cultivated lettuce. Differentiations in bacterial community structure translated into differences in microbiome function, including an increase in avirulence factors contributing to the increased susceptibility of cultivated lettuce to pathogens, indicating that changes in the microbiome of domesticated lettuce may translate into a reduction in fitness. Wild lettuce accessions have been identified as sources for resistance to bacterial, fungal and viral pathogens and have been exploited in disease-resistant germplasm development (Lebeda *et al.*, 2014). Wild lettuce is also known to differ significantly in secondary metabolites, with an increase in phenolic compounds observed compared to cultivated lettuce (Damerum *et al.*, 2015), which aside from leaf surface morphology may impact on microbial diversity.

The core lettuce bacterial microbiome determined from 171 RILs as defined here, was comprised largely of Proteobacteria in the class Gammaproteobacteria; *Pseudomonas*, *Alkanindiges*, *Acinetobacter*, Alphaproteobacteria; *Sphingomonas*, and Betaproteobacteria; *Massilia*, all previously determined to commonly exist in the phyllosphere of diverse plant species (Thapa & Prasanna, 2018). *Buchnera*, an unculturable aphid endosymbiont representing on average ~18.4% of the relative abundance across the population, has previously been identified at similar relative abundances on spinach leaves (Burch *et al.*, 2016). Total species richness (observed OTUs), evenness (Pielou’s index) and Shannon indexes were comparable to other leaf bacterial microbiomes reported (Lopez-Velasco *et al.*, 2011; Roman-Reyna *et al.*, 2019). The most abundantly observed taxa (*Pseudomonas* and *Buchnera* spp.) were negatively correlated with most α and β diversity measures, indicating that their presence was associated with reduced phyllosphere diversity. Interestingly, the most abundant *Buchnera* or *Pseudomonas* taxa demonstrated few negative interactions with other relatively low abundance taxa in network analyses (Table S9), suggesting this negative association may be passive, through resource or space competition, as opposed to active, caused by attack. Since *Pseudomonas* spp. are important spoilage organisms, this finding is worthy of further consideration through future manipulation experiments within synthetic communities, to test the role of diversity in determining the abundance of specific *Pseudomonas* species. The lettuce bacterial community observed in >5% of the RILs was composed of a small number (19) of abundant taxa (>1% abundance) and a larger number (1,427 taxa) at low abundance (<1% abundance). Potential human pathogens including *E. coli*, although sparsely identified (~11% of samples) and at low abundance (<1%), were nevertheless present, alongside several other human pathogenic bacteria. Though *E. coli* is not well-adapted to epiphytic survival, *E. coli* inoculated onto lettuce leaves has been shown to persist and still be observed after 4 weeks (Moyne *et al.*, 2011). Our data may provide insights into particular plant genotypes where these human pathogens remain, which are important resources for future investigation, alongside a range of genotypes where *E. coli* was not present.

Co-occurrence was observed more frequently than co-exclusion, but overall, taxa demonstrated weak interactions as reflected by the low network modularity observed in cooccurrence analyses. For the first time in lettuce, keystone species were detected which have an important influence on overall bacterial community structure. These species were present at low abundance on the majority (>99%) of leaves. Keystone taxa included a *Terrabacter* spp., present on all RIL samples and three taxa in the Sphingomonadaceae family. *Terrabacter* spp. have been reported in agricultural soils (Sugiyama *et al.*, 2017) and on the surface of *Eucalyptus* leaves (Miguel *et al.*, 2016) but have not been previously identified in the lettuce phyllosphere. *Sphingomonas* spp. are commensals which have been identified as keystone species in *Arabidopsis* (Carlström *et al.*, 2019) and have been demonstrated to promote plant growth (Khan *et al.*, 2014). Members of the *Sphingomonas* genus have been shown to supress the pathogenicity of *Pseudomonas syringae* (Innerebner *et al.*, 2011) and Vogel *et al.* (2016) demonstrated that inoculating *Arabidopsis* with *Sphingomonas melonis* elicited an immune-like transcriptional response, hypothesised to prime the plant for subsequent response to pathogenic *P. syringae*. Elucidating the molecular mechanisms of such interactions between microbiota and plant in this food crop will be crucial in determining how they contribute to plant health and fitness and how they can be engineered in future.

Leaf characteristics including total leaf area, abaxial surface stomatal size and hydrophobicity and adaxial surface epidermal cell number, area and stomatal density were identified to influence overall community structure. Reduced abaxial stomata pore size was negatively correlated with all α diversity measures (r=0.2, P<0.05), had a significant influence on β diversity distance (PERMANOVA, P<0.05) and on PCA of bacterial community structure (R^2^=0.05-0.08, P<0.01). Larger stomata are known to aid the entry of endophytic bacteria into the leaf, with wider stomata pores leading to increased bacterial infiltration into the leaf by chemotaxis towards photosynthetic products within the leaf (Ranjbaran *et al.*, 2020). Surfaces with wider stomata and a more hydrophilic surface (increased contact angle) have also been shown to enhance infiltration of bacteria through stomatal pores present on the surface in droplet suspensions (Ranjbaran & Datta, 2019). Three *Pseudomonas* taxa were identified at reduced abundance on leaves with smaller stomata and two *Pseudomonas* taxa were identified at a greater abundance on leaves with reduced size, perhaps indicating interspecies competition in the *Pseudomonas* genus between endo- and epiphytic species. Indeed *Pseudomonas* has been identified to inhabit both compartments (Karasov *et al.*, 2018). The bacterial microbiome of leaves with smaller stomata also had reduction in the abundance of genes functioning in plant-pathogen interactions, chemotaxis and quorum sensing, suggesting stomatal size as a key trait for future plant breeding, which could incidentally have co-benefits with respect to water-use efficiency, although this remains to be tested (Bertolino *et al.*, 2019). Leaf size was also important in influencing the bacterial community and leaves with a smaller area harboured a microbiome with a reduced abundance of genes functioning in biofilm formation. Leaves with a low contact angle, indicating a more hydrophilic surface, had a reduced abundance of a *Sphingomonas* spp., likely due to differences in mobility on these surfaces.

QTL for bacterial abundance, for the first time in lettuce, and leaf trait phenotypes were mapped in a commercial field environment, identifying novel host loci underlying these traits. QTL for both community-level properties (diversity metrics) and individual taxa were identified, suggesting both are influenced by plant genetics. QTL for α and β diversity metrics often mapped to the same position in the lettuce genome, identifying regions important in shaping whole community structure and likely to be significant in determining fitness in lettuce. QTL for bacterial taxa were often clustered and mapped to the same position as QTL for host phenotypes including stomata size and leaf area, indicating that regions of the host genome controlling leaf surface traits can influence the abundance of multiple taxa simultaneously. A single QTL was identified for each of the top five most abundant taxa, including *Buchnera*, *Pseudomonas* and *Alkanindiges* spp., indicating that host genetics had limited influence on the most abundant taxa, which are perhaps more influenced by the timing and order in which taxa arrive on the leaf surface.

As previously observed (Horton *et al.*, 2014), there was an abundance of cell wall-associated genes within genomic regions and enrichment of cell wall-associated GO terms where ASVs mapped. This is unsurprising given the role of the cell wall as the first obstacle microorganisms encounter and is a target for pathogen attack by cell wall degradation. Hormone signalling, defence response and response to stress were also significantly overrepresented, with auxin- and ethylene-responsive genes identified within all candidate regions investigated. Phytohormones are widely recognised to have crucial roles in plant defence response to biotic stressors, regulating the colonisation of both beneficial and pathogenic microbes (Eichmann *et al.*, 2021). The ethylene-mediated pathway is associated with plant response to necrotrophic pathogens and the inhibition of leaf growth and cell division in response to abiotic stress (Dubois *et al.*, 2018). Ethylene and auxin have been demonstrated to have antagonistic effects on abscisic acid-induced stomata closure (Tanaka *et al.*, 2006). Endogenous application of auxin (indole acetic acid) to *Streptomyces* strains isolated from *Arabidopsis* roots elicited antibacterial activity towards *Escherichia coli* and *Bacillus subtilis*, providing evidence for a “cry for help” mechanism, whereby plants produce phytohormones to stimulate antibiotic production of their inhabiting microbiota (van der Meij *et al.*, 2018; Eichmann *et al.*, 2021). Leucine-rich repeat domain- and lectin domain-containing proteins are important in the perception of microbe-/pathogen- and damage-associated molecular patterns (MAMPs/PAMPs/DAMPs), recognising carbohydrate structures derived from microbes (MAMPs or PAMPs) or host carbohydrates, like those comprising the cell wall, which result from pathogen attack (DAMPs), triggering an immune response (Lannoo & Van Damme, 2014). Taken together, our results indicate that host influence on the lettuce bacterial microbiome is shaped by genes regulating stomata morphology, cell wall biosynthesis and maintenance, hormone-induced signalling responses and pathogen detection receptors and these areas of the genome provide important targets to inform future crosses to enable hypothesis testing and strategies for molecular breeding. Future research will focus on enhancing yields of beneficial microbes (investigating two areas of linkage group 1, where several keystone taxa were mapped) and reducing pathogenic and spoilage microorganisms (considering linkage groups 4, 5 and 6, where key taxa were mapped in several QTL hotspots). Thus, we have taken the first steps in unravelling the genetic basis of the ‘good’ and ‘bad’ lettuce microbiome, providing new knowledge and understanding of the complexities of this ecosystem. This now requires further manipulation and experimentation to test the importance of these genomic regions for improved lettuce.

## CONCLUSION

This study provides the first QTL for bacteria abundance, alongside leaf host trait QTL, in a biparental mapping population of an important food crop. We identified the genetic architecture of key leaf surface characteristics, including stomata and epidermal cell size and leaf area to be important host traits effecting the phyllosphere bacterial microbiome, including for spoilage and human pathogenic bacteria. Through co-occurrence network analyses, we identified keystone taxa which have a high influence on overall bacterial community structure. Further studies will involve teasing apart the environmental influence of host effects in multi-site studies and in testing our findings in specific lettuce genotypes, to assess the potential of using microbiome engineering to develop lettuce with improved food safety and reduced spoilage characteristics.

## Supporting information

Figure S1D part 4

Figure S1D part 3

Figure S1D part 2

Figure S1D part 1

Figure S1C part 2

Figure S1C part 1

## ACKNOWLEDGEMENTS

This research was funded by the Biological and Biotechnological Sciences Research Council (BBSRC) through a PhD award to ECA and by Vitacress Salads Ltd. Funding was also provided by The Californian Leafy Greens Research Board, by the University of California, Davis and by the John B. Orr Endowed Chair in Environmental Plant Sciences, held by GT. We thank the laboratory of Richard W. Michelmore at the University of California, Davis for providing the core RIL lettuce mapping population and associated genomic resources. We thank S. Milner, B. Valdes and I. Greffet for field assistance.

## AUTHOR CONTRIBUTIONS

AD conducted the data analysis, drafted the manuscript and assisted in data collection. ECA conducted the field trial, sample analysis and data collection and VB assisted with data analysis. GT was PI and conceived and led the project and all authors commented on the manuscript.

## REFERENCES

Alegbeleye OO, Singleton I, Sant’Ana AS. 2018. Sources and contamination routes of microbial pathogens to fresh produce during field cultivation: A review. Food Microbiology 73: 177–208.

Barkan A, Small I. 2014. Pentatricopeptide repeat proteins in plants. Annual Review of Plant Biology 65: 415–442.

Beilsmith K, Thoen MPM, Brachi B, Gloss AD, Khan MH, Bergelson J. 2019. 3kkiiGenome-wide association studies on the phyllosphere microbiome: Embracing complexity in host– microbe interactions. Plant Journal 97: 164–181.

Bennett SD, Sodha S V, Ayers TL, Lynch MF, Gould LH, Tauxe R V. 2018. Produce-associated foodborne disease outbreaks, USA, 1998–2013. Epidemiology and Infection 146: 1397–1406.

Bertolino LT, Caine RS, Gray JE. 2019. Impact of stomatal density and morphology on water-use efficiency in a changing world. Frontiers in Plant Science 10.

Bokulich NA, Kaehler BD, Rideout JR, Dillon M, Bolyen E, Knight R, Huttley GA, Caporaso JG. 2018. Optimizing taxonomic classification of marker-gene amplicon sequences with QIIME 2 ’s q2-feature-classifier plugin. : 1–17.

Bolyen E, Rideout JR, Dillon MR, Bokulich NA, Abnet CC, Al-Ghalith GA, Alexander H, Alm EJ, Arumugam M, Asnicar F, et al. 2019. Reproducible, interactive, scalable and extensible microbiome data science using QIIME 2. Nature Biotechnology 37: 852–857.

Brandl MT. 2008. Plant lesions promote the rapid multiplication of Escherichia coli O157:H7 on postharvest lettuce. Applied and Environmental Microbiology 74: 5285–5289.

Broman KW. 2003. Mapping quantitative trait loci in the case of a spike in the phenotype distribution. Genetics 163: 1169–1175.

Broman KW. 2009. A guide to QTL mapping with R / qtl.

Burch AY, Do PT, Sbodio A, Suslow T V., Lindow SE. 2016. High-level culturability of epiphytic bacteria and frequency of biosurfactant producers on leaves. Applied and Environmental Microbiology 82: 5997–6009.

Callahan BJ, McMurdie PJ, Rosen MJ, Han AW, Johnson AJA, Holmes SP. 2016. DADA2: High-resolution sample inference from Illumina amplicon data. Nature Methods 13: 581–583.

Cardinale M, Grube M, Erlacher A, Quehenberger J, Berg G. 2015. Bacterial networks and co-occurrence relationships in the lettuce root microbiota. Environmental Microbiology 17: 239–252.

Carlström CI, Field CM, Bortfeld-miller M, Müller B, Sunagawa S, Vorholt JA. 2019. Synthetic microbiota reveal priority effects and keystone strains in the Arabidopsis phyllosphere. Nature Ecology & Evolution 3: 1445–1454.

Castillo AI, Nelson ADL, Haug-baltzell AK. 2018. A tutorial of diverse genome analysis tools found in the CoGe web-platform using Plasmodium spp. as a model. Database 2018: 1–16.

Copeland JK, Yuan L, Layeghifard M, Wang PW, Guttman DS. 2015. Seasonal community succession of the phyllosphere microbiome. Molecular Plant-Microbe Interactions 28: 274–285.

Csardi G, Nepusz T. 2006. The igraph software package for complex network research. InterJournal Complex Sy: 1–9.

Damerum A, Selmes SL, Biggi GF, Clarkson GJ, Rothwell SD, Truco MJ, Michelmore RW, Hancock RD, Shellcock C, Chapman M a, et al. 2015. Elucidating the genetic basis of antioxidant status in lettuce (Lactuca sativa). Horticulture Research 2: 1–13.

Damerum A, Smith HK, Clarkson GJJ, Truco MJ, Michelmore RW, Taylor G. 2021. The genetic basis of water-use efficiency and yield in lettuce. BMC Plant Biology 21: 1–14.

Douglas G, Maffei V, Zaneveld J, Yurgel S, Brown J, Taylor C, Huttenhower C, Langille M. 2020. PICRUSt2 for prediction of metagenome functions. Nat Biotechnology 38: 669–688.

Doyle JJ, Doyle JL. 1987. A rapid DNA isolation procedure for small quantities of fresh leaf tissue. Phytochemical Bulletin 19: 11–15.

Dubois M, Van den Broeck L, Inzé D. 2018. The Pivotal Role of Ethylene in Plant Growth. Trends in Plant Science 23: 311–323.

Dufayard JF, Bettembourg M, Fischer I, Droc G, Guiderdoni E, Périn C, Chantret N, Diévart A. 2017. New insights on Leucine-Rich repeats receptor-like kinase orthologous relationships in angiosperms. Frontiers in Plant Science 8: 1–18.

Eichmann R, Richards L, Schäfer P. 2021. Hormones as go-betweens in plant microbiome assembly. Plant Journal 105: 518–541.

Ge SX, Jung D, Jung D, Yao R. 2020. ShinyGO: A graphical gene-set enrichment tool for animals and plants. Bioinformatics 36: 2628–2629.

Glöckner FO, Yilmaz P, Quast C, Gerken J, Beccati A, Ciuprina A, Bruns G, Yarza P, Peplies J, Westram R, et al. 2017. 25 years of serving the community with ribosomal RNA gene reference databases and tools. Journal of Biotechnology 261: 169–176.

Herman KM, Hall AJ, Gould LJ. 2015. Outbreaks attributed to fresh leafy vegetables, United States, 1973–2012. Epidemiology and Infection 143: 3011–3021.

Highmore CJ, Warner JC, Rothwell SD, Wilks SA, Keevil CW. 2018. Viable-but-Nonculturable Listeria monocytogenes and Salmonella enterica Serovar Thompson Induced by Chlorine Stress Remain Infectious Callum. mBio 9: e00540–18.

Horton MW, Bodenhausen N, Beilsmith K, Meng D, Muegge BD, Subramanian S, Vetter MM, Vilhjálmsson BJ, Nordborg M, Gordon JI, et al. 2014. Genome-wide association study of Arabidopsis thaliana leaf microbial community. Nature Communications 5: 1–7.

Hunter PJ, Shaw RK, Berger CN, Frankel G, Pink D, Hand P. 2015. Older leaves of lettuce (Lactuca spp.) support higher levels of Salmonella enterica ser. Senftenberg attachment and show greater variation between plant accessions than do younger leaves. FEMS Microbiology Letters 362: 1–7.

Innerebner G, Knief C, Vorholt JA. 2011. Protection of Arabidopsis thaliana against leaf-pathogenic Pseudomonas syringae by Sphingomonas strains in a controlled model system. Applied and Environmental Microbiology 77: 3202–3210.

Jacob C, Melotto M. 2020. Human Pathogen Colonization of Lettuce Dependent Upon Plant Genotype and Defense Response Activation. Frontiers in Plant Science 10: 1–17.

Karasov TL, Almario J, Friedemann C, Ding W, Giolai M, Heavens D, Kersten S, Lundberg DS, Neumann M, Regalado J, et al. 2018. Arabidopsis thaliana and Pseudomonas Pathogens Exhibit Stable Associations over Evolutionary Timescales. Cell Host and Microbe 24: 168–179.e4.

Kembel SW, O’Connor TK, Arnold HK, Hubbell SP, Wright SJ, Green JL. 2014. Relationships between phyllosphere bacterial communities and plant functional traits in a neotropical forest. Proceedings of the National Academy of Sciences of the United States of America 111: 13715–13720.

Khan AL, Waqas M, Kang SM, Al-Harrasi A, Hussain J, Al-Rawahi A, Al-Khiziri S, Ullah I, Ali L, Jung HY, et al. 2014. Bacterial endophyte Sphingomonas sp. LK11 produces gibberellins and IAA and promotes tomato plant growth. Journal of Microbiology 52: 689–695.

Koskella B. 2020. The phyllosphere. Current Biology 30: R1143–R1146.

Krzywinski M, Schein J, Birol I, Connors J, Gascoyne R, Horsman D, Jones SJ, Marra MA. 2009. Circos: An information aesthetic for comparative genomics. Genome Research 19: 1639–1645.

Kurtz ZD, Müller CL, Miraldi ER, Littman DR, Blaser MJ, Bonneau RA. 2015. Sparse and Compositionally Robust Inference of Microbial Ecological Networks. PLoS Computational Biology 11: 1–25.

Laforest-Lapointe I, Paquette A, Messier C, Kembel SW. 2017. Leaf bacterial diversity mediates plant diversity and ecosystem function relationships. Nature 546: 145–147.

Lamour G, Hamraoui A, Buvailo A, Xing Y, Keuleyan S, Prakash V, Eftekhari-Bafrooei A, Borguet E. 2010. Contact angle measurements using a simplified experimental setup. Journal of Chemical Education 87: 1403–1407.

Lannoo N, Van Damme EJM. 2014. Lectin domains at the frontiers of plant defense. Frontiers in Plant Science 5: 1–16.

Lebeda A, Křístková E, Kitner M, Mieslerová B, Jemelková M, Pink D a. C. 2014. Wild Lactuca species, their genetic diversity, resistance to diseases and pests, and exploitation in lettuce breeding. European Journal of Plant Pathology 138: 597–640.

Liu H, Brettell LE, Singh B. 2020. Linking the Phyllosphere Microbiome to Plant Health. Trends in Plant Science 25: 841–844.

López-Gálvez F, Gil MI, Truchado P, Selma M V., Allende A. 2010. Cross-contamination of fresh-cut lettuce after a short-term exposure during pre-washing cannot be controlled after subsequent washing with chlorine dioxide or sodium hypochlorite. Food Microbiology 27: 199–204.

Lopez-Velasco G, Welbaum GE, Boyer RR, Mane SP, Ponder MA. 2011. Changes in spinach phylloepiphytic bacteria communities following minimal processing and refrigerated storage described using pyrosequencing of 16S rRNA amplicons. Journal of Applied Microbiology 110: 1203–1214.

Love MI, Huber W, Anders S. 2014. Moderated estimation of fold change and dispersion for RNA-seq data with DESeq2. Genome Biology 15: 550.

Mcmurdie PJ, Holmes S. 2013. phyloseq : An R Package for Reproducible Interactive Analysis and Graphics of Microbiome Census Data. 8.

van der Meij A, Willemse J, Schneijderberg MA, Geurts R, Raaijmakers JM, van Wezel GP. 2018. Inter- and intracellular colonization of Arabidopsis roots by endophytic actinobacteria and the impact of plant hormones on their antimicrobial activity. Antonie van Leeuwenhoek, International Journal of General and Molecular Microbiology 111: 679–690.

Miguel PSB, de Oliveira MNV, Delvaux JC, de Jesus GL, Borges AC, Tótola MR, Neves JCL, Costa MD. 2016. Diversity and distribution of the endophytic bacterial community at different stages of Eucalyptus growth. Antonie van Leeuwenhoek 109: 755–771.

Moyne AL, Sudarshana MR, Blessington T, Koike ST, Cahn MD, Harris LJ. 2011. Fate of Escherichia coli O157:H7 in field-inoculated lettuce. Food Microbiology 28: 1417–1425.

Oksanen J, Blanchet FG, Friendly M, Kindt R, Legendre P, McGlinn D, Minchin PR, O’Hara RB, Simpson GL, Solymos P, et al. 2020. vegan: Community Ecology Package. : 2.5–7.

Price MN, Dehal PS, Arkin AP. 2010. FastTree 2 – Approximately Maximum-Likelihood Trees for Large Alignments. PloS ONE 5: e9490.

R Core Team. 2017. R: A Language and Environment for Statistical Computing. 7: 275–286.

Ragaert P, Devlieghere F, Debevere J. 2007. Role of microbiological and physiological spoilage mechanisms during storage of minimally processed vegetables. Postharvest Biology and Technology 44: 185–194.

Ranjbaran M, Datta AK. 2019. Retention and infiltration of bacteria on a plant leaf driven by surface water evaporation. Physics of Fluids 31.

Ranjbaran M, Solhtalab M, Datta AK. 2020. Mechanistic modeling of light-induced chemotactic infiltration of bacteria into leaf stomata. PLoS Computational Biology 16: 1–23.

Rognes T, Flouri T, Nichols B, Quince C, Mahé F. 2016. VSEARCH: A versatile open source tool for metagenomics. PeerJ 2016: 1–22.

Roman-Reyna V, Pinili D, Borja FN, Quibod IL, Groen SC, Mulyaningsih ES, Rachmat A, Slamet-Loedin IH, Alexandrov N, Mauleon R, et al. 2019. The rice leaf microbiome has a conserved community structure controlled by complex host-microbe interactions. bioRxiv.

Sathoff AE, Sathoff AE, Lewenza S, Lewenza S, Samac DA, Samac DA. 2020. Plant defensin antibacterial mode of action against Pseudomonas species. BMC Microbiology 20: 1–11.

Schneider C a, Rasband WS, Eliceiri KW. 2012. NIH Image to ImageJ: 25 years of image analysis. Nature Methods 9: 671–675.

Sugiyama A, Unno Y, Ono U, Yoshikawa E, Suzuki H, Minamisawa K, Yazaki K. 2017. Assessment of bacterial communities of black soybean grown in fields. Communicative and Integrative Biology 10: 1–8.

Tanaka Y, Sano T, Tamaoki M, Nakajima N, Kondo N, Hasezawa S. 2006. Cytokinin and auxin inhibit abscisic acid-induced stomatal closure by enhancing ethylene production in Arabidopsis. Journal of Experimental Botany 57: 2259–2266.

Thapa S, Prasanna R. 2018. Prospecting the characteristics and significance of the phyllosphere microbiome. Annals of Microbiology 68: 229–245.

Trivedi P, Leach JE, Tringe SG, Sa T, Singh BK. 2020. Plant–microbiome interactions: from community assembly to plant health. Nature Reviews Microbiology 18: 607–621.

Truco MJ, Antonise R, Lavelle D, Ochoa O, Kozik A, Witsenboer H, Fort SB, Jeuken MJW, Kesseli R V, Lindhout P, et al. 2007. A high-density, integrated genetic linkage map of lettuce (Lactuca spp.). Theoretical and Applied Genetics 115: 735–46.

Truco MJ, Ashrafi H, Kozik A, van Leeuwen H, Bowers J, Reyes Chin Wo S, Stoffel K, Xu H, Hill T, Van Deynze A, et al. 2013. An Ultra High-Density, Transcript-Based, Genetic Map of Lettuce. G3: Genes | Genomes | Genetics 3: 617–631.

USDA. 2020. Vegetables 2019 summary.

Villanueva RAM, Chen ZJ. 2019. ggplot2: Elegant Graphics for Data Analysis (2nd ed.). Measurement: Interdisciplinary Research and Perspectives 17: 160–167.

Vogel C, Bodenhausen N, Gruissem W, Vorholt JA. 2016. The Arabidopsis leaf transcriptome reveals distinct but also overlapping responses to colonization by phyllosphere commensals and pathogen infection with impact on plant health. The New phytologist 212: 192–207.

Wei T, Simko V. 2021. R package ‘corrplot’: Visualization of a Correlation Matrix. : (Version 0.88).

Williams TR, Moyne AL, Harris LJ, Marco ML. 2013. Season, Irrigation, Leaf Age, and Escherichia coli Inoculation Influence the Bacterial Diversity in the Lettuce Phyllosphere. PLoS ONE 8: 1–14.

Zeng W, Melotto M, He SY. 2010. Plant stomata: A checkpoint of host immunity and pathogen virulence. Current Opinion in Biotechnology 21: 599–603.

Zhang FZ, Wagstaff C, Rae AM, Sihota AK, Keevil CW, Rothwell SD, Clarkson GJJ, Michelmore RW, Truco MJ, Dixon MS, et al. 2007. QTLs for shelf life in lettuce co-locate with those for leaf biophysical properties but not with those for leaf developmental traits. Journal of Experimental Botany 58: 1433–1449.

