## Supplementary figures and images for "Good and bad lettuce leaf microbes? Unravelling the genetic architecture of the microbiome to inform plant breeding for enhanced food safety and reduced food waste"

### Figure S1A

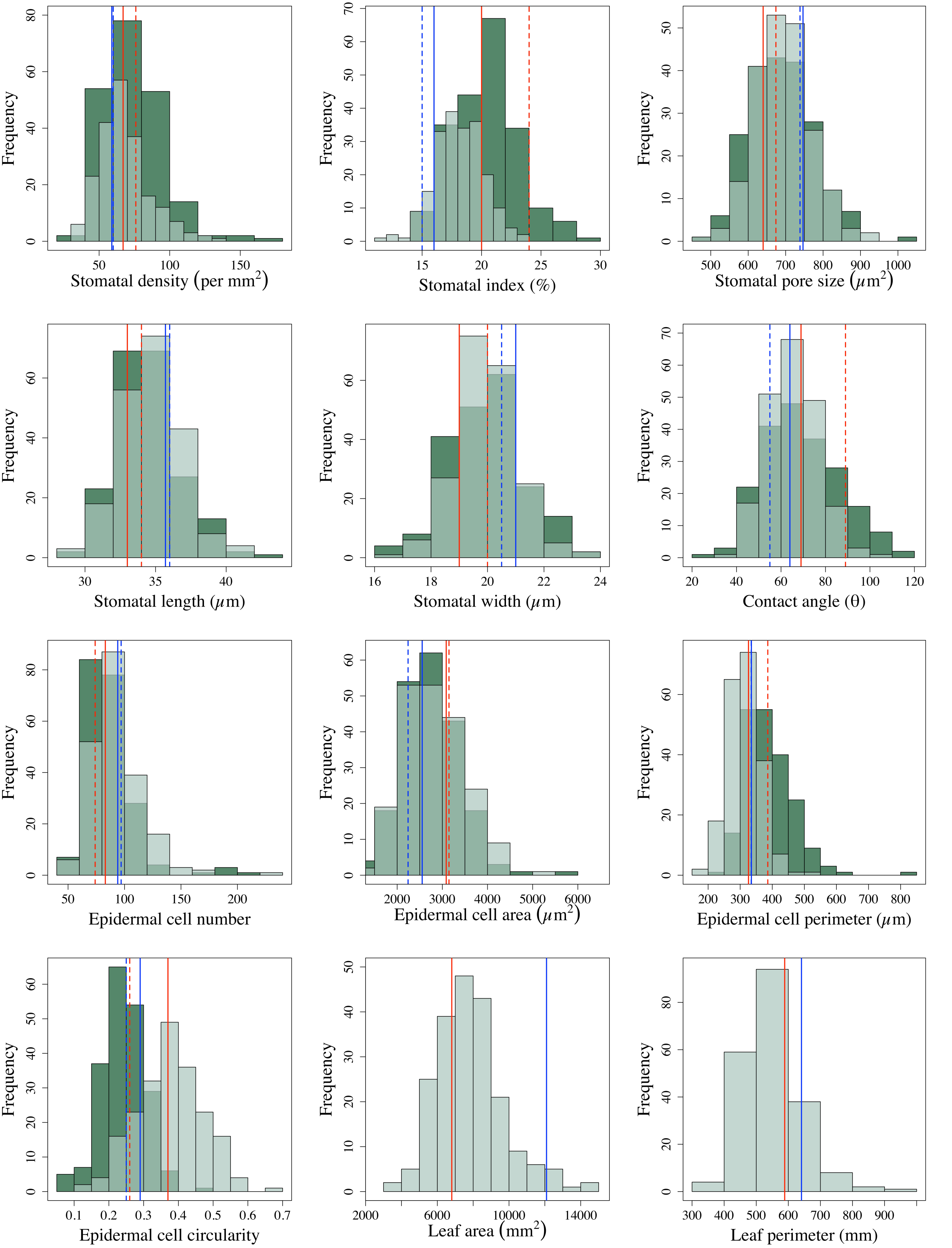

### Figure S1B

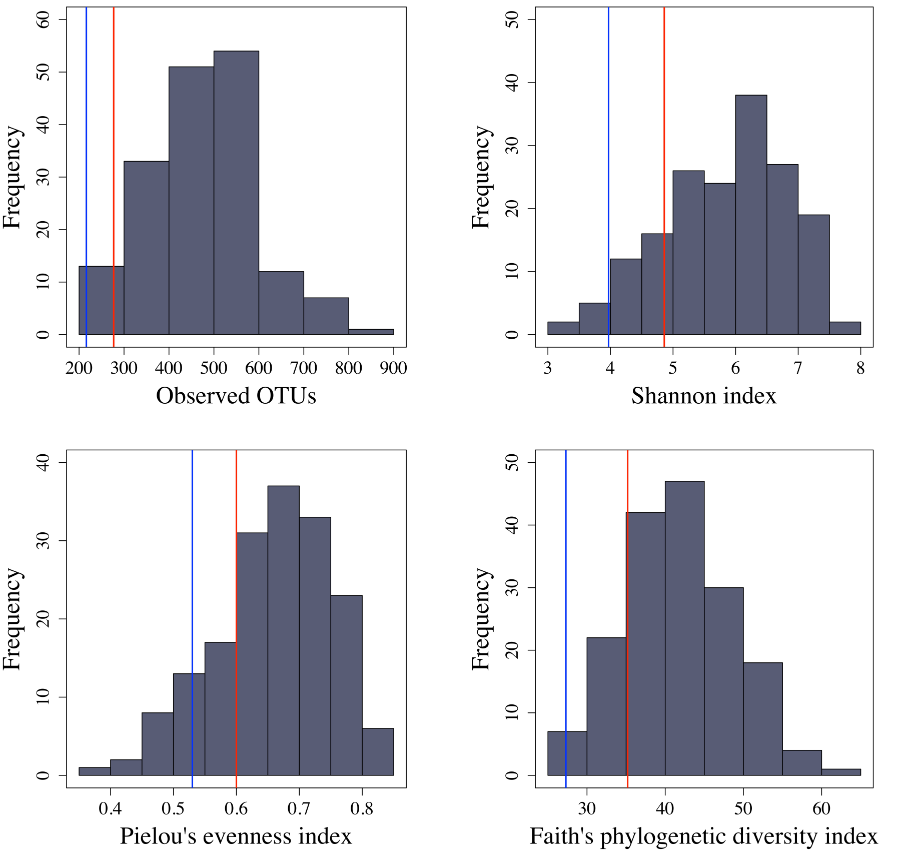

### Figure S1C part 1

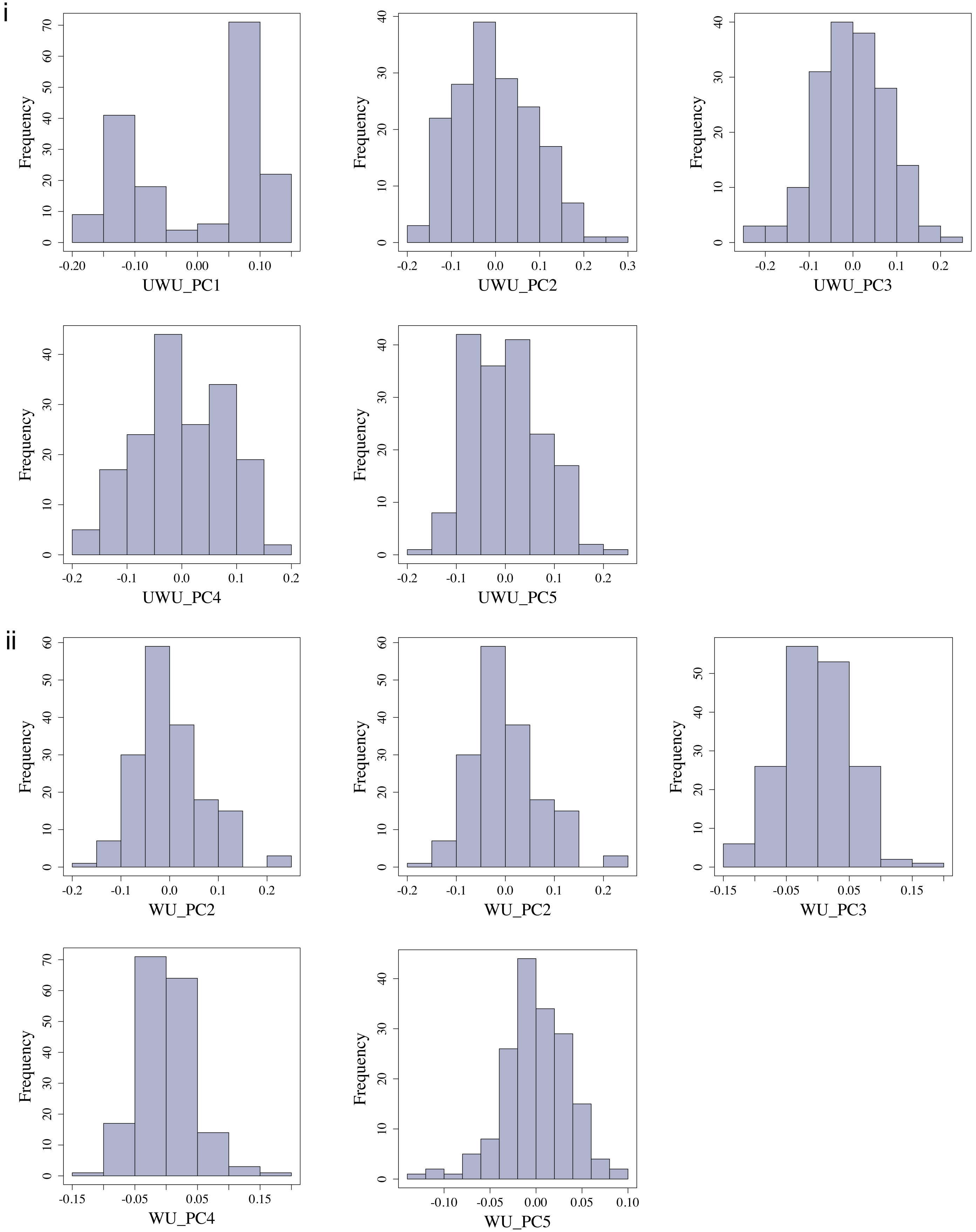

### Figure S1C part 2

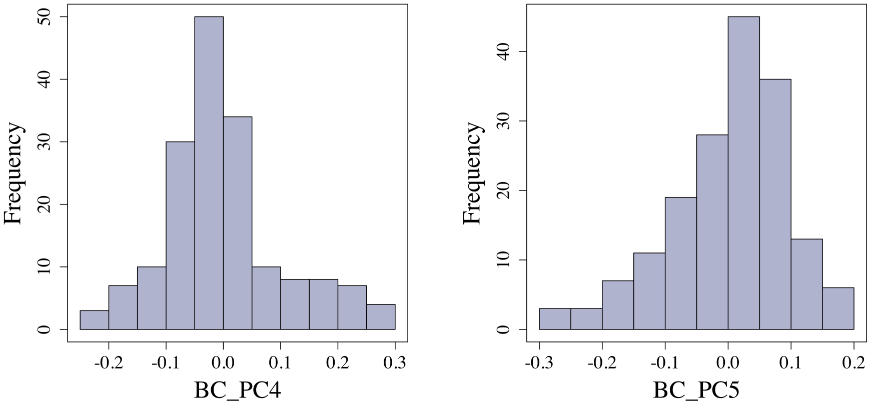

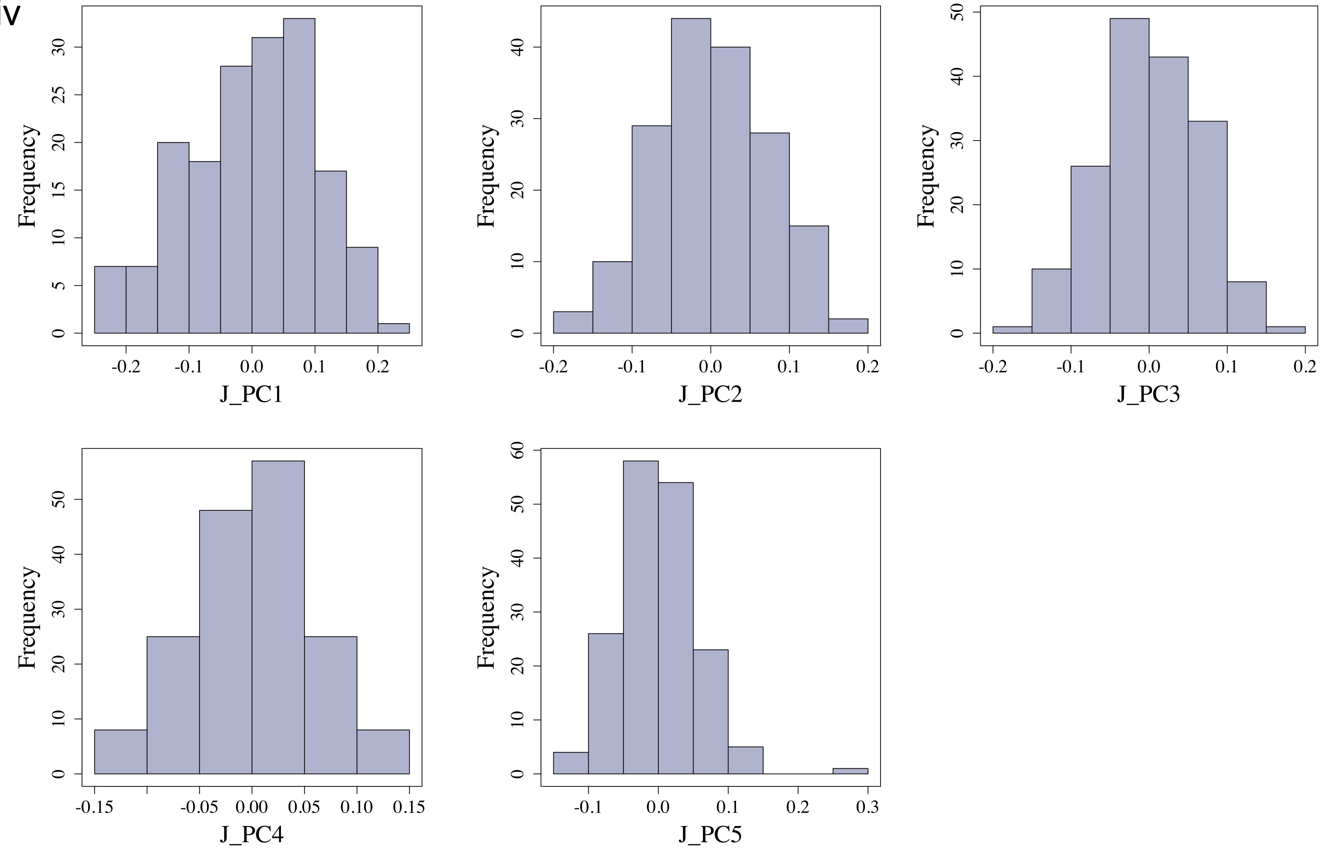

### Figure S1D part 1

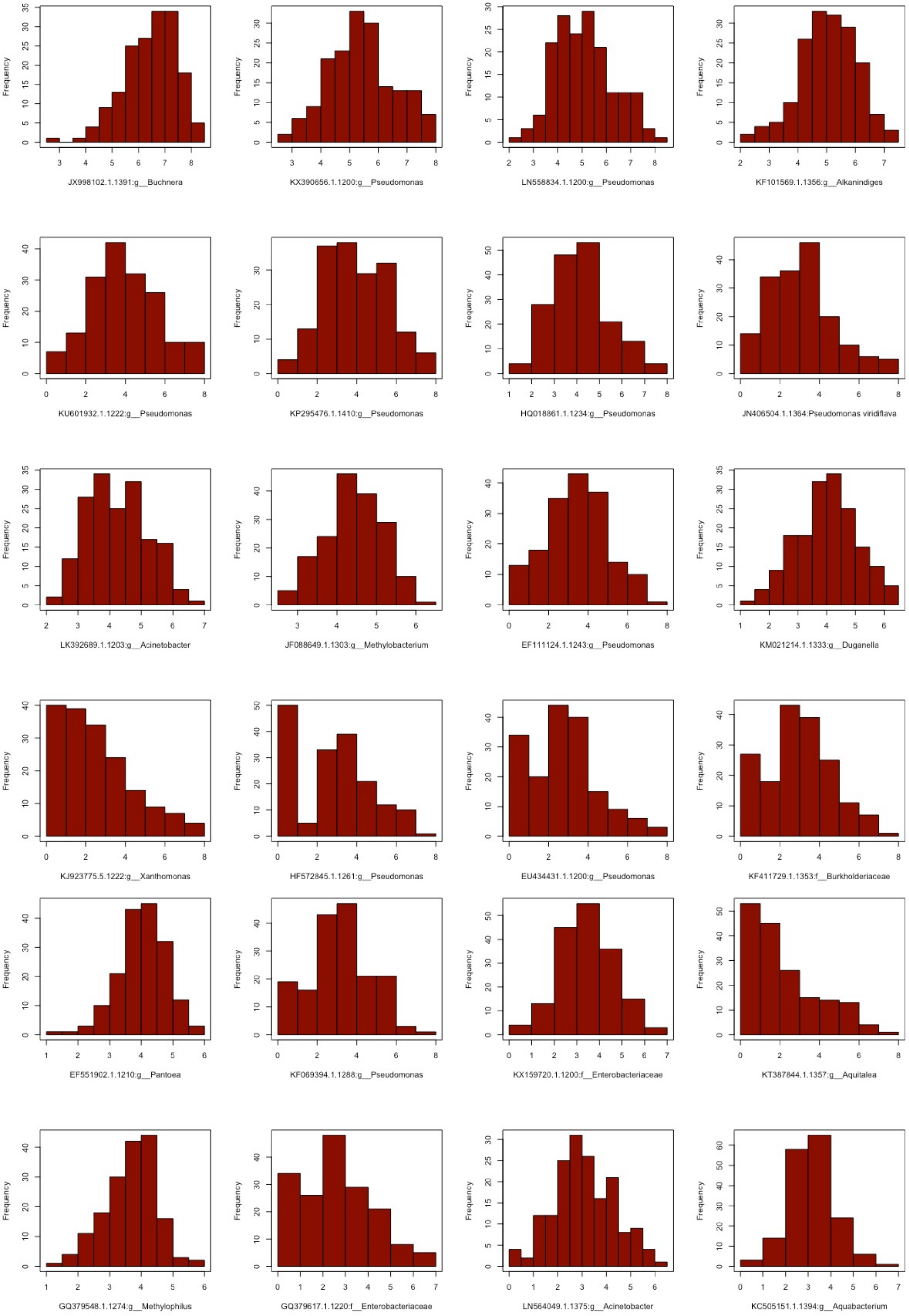

### Figure S1D part 2

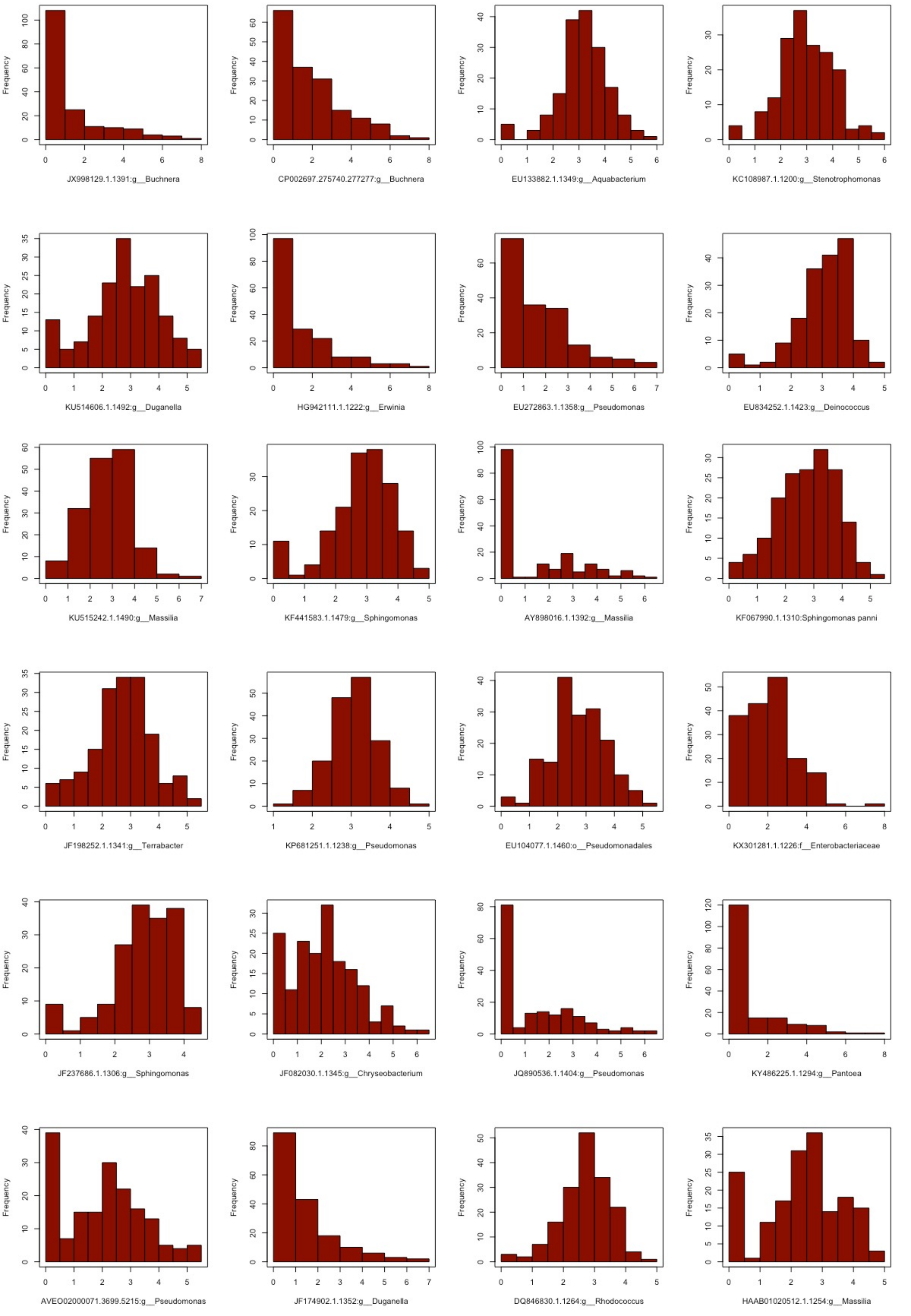

### Figure S1D part 3

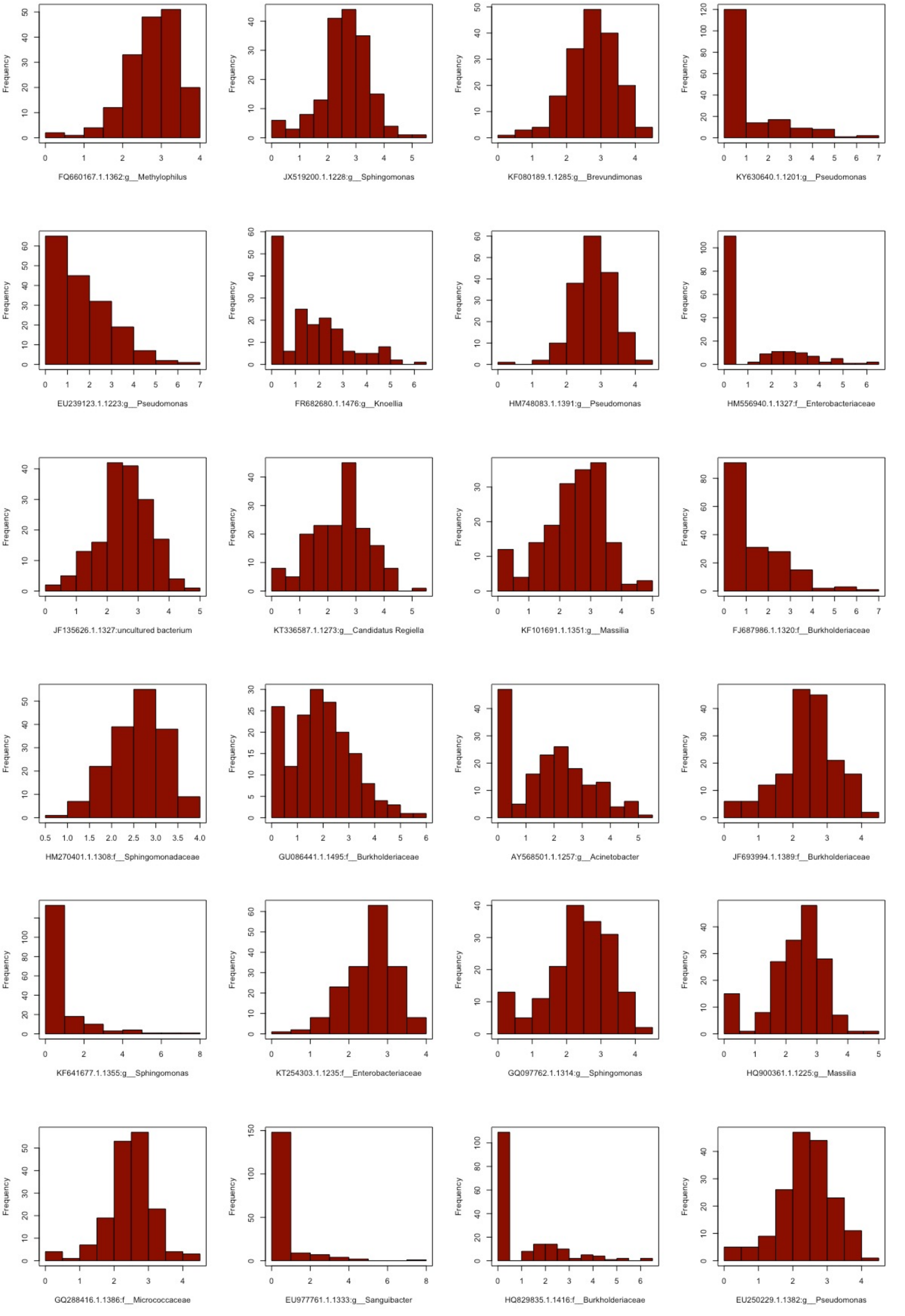

### Figure S1D part 4

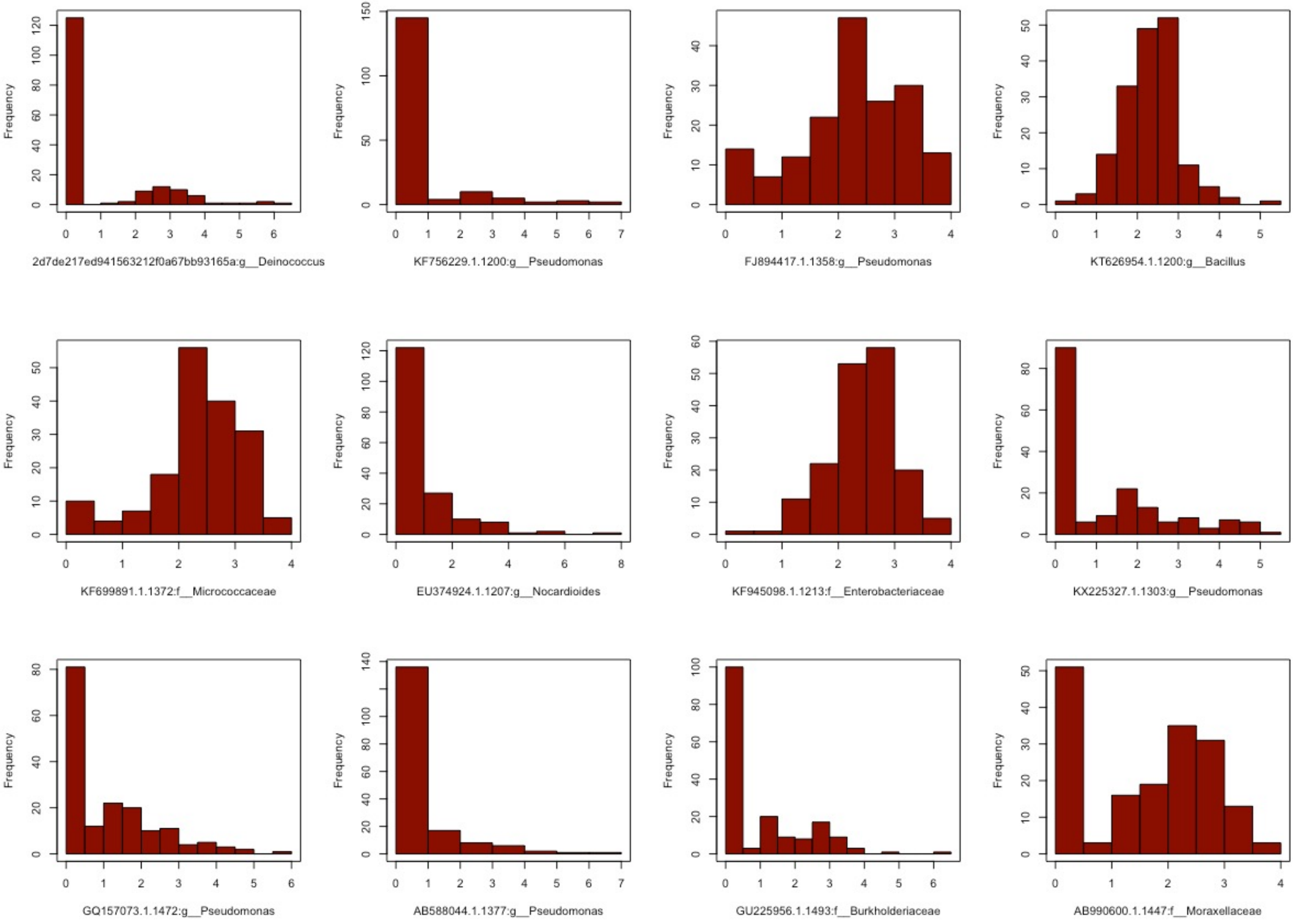
